# Ancestral Haplotype Reconstruction in Endogamous Populations using Identity-By-Descent

**DOI:** 10.1101/2020.01.15.908459

**Authors:** Kelly Finke, Michael Kourakos, Gabriela Brown, Huyen Trang Dang, Shi Jie Samuel Tan, Yuval B. Simons, Shweta Ramdas, Alejandro A. Schäffer, Rachel L. Kember, Maja Bućan, Sara Mathieson

## Abstract

In this work we develop a novel algorithm for reconstructing the genomes of ancestral individuals, given genotype or sequence data from contemporary individuals and an extended pedigree of family relationships. A pedigree with complete genomes for every individual enables the study of allele frequency dynamics and haplotype diversity across generations, including deviations from neutrality such as transmission distortion. When studying heritable diseases, ancestral haplotypes can be used to augment genome-wide association studies and track disease inheritance patterns. The building blocks of our reconstruction algorithm are segments of Identity-By-Descent (IBD) shared between two or more genotyped individuals. The method alternates between identifying a source for each IBD segment and assembling IBD segments placed within each ancestral individual. Unlike previous approaches, our method is able to accommodate complex pedigree structures with hundreds of individuals genotyped at millions of SNPs.

We apply our method to an Old Order Amish pedigree from Lancaster, Pennsylvania, whose founders came to the United States from Europe during the early 18th century. The pedigree includes 1338 individuals from the past 10 generations, 394 with genotype data. The motivation for reconstruction is to understand the genetic basis of diseases segregating in the family through tracking haplotype transmission over time. Using our algorithm thread, we are able to reconstruct an average of 224 ancestral individuals per chromosome. For these ancestral individuals, on average we reconstruct 79% of their haplotypes. We also identify a region on chromosome 16 that is difficult to reconstruct – we find that this region harbors a short Amish-specific copy number variation and the gene *HYDIN*. `thread` was developed for endogamous populations, but can be applied to any extensive pedigree with the recent generations genotyped. We anticipate that this type of practical ancestral reconstruction will become more common and necessary to understand rare and complex heritable diseases in extended families.

**Author summary:** When analyzing complex heritable traits, it is often useful to have genomic data from many generations of an extended family, to increase the amount of information available for statistical inference. However, we typically only have genomic data from the recent generations of a pedigree, as ancestral individuals are deceased. In this work we present an algorithm, called `thread`, for reconstructing the genomes of ancestral individuals, given a complex pedigree and genomic data from the recent generations. Previous approaches have not been able to accommodate large datasets (both in terms of sites and individuals), made simplifying assumptions about pedigree structure, or did not tie reconstructed sequences back to specific individuals. We apply `thread` to a complex Old Order Amish pedigree of 1338 individuals, 394 with genotype data.

## Introduction

Pedigree structures and associated genetic data provide a wealth of information for studying recent evolution. Nuclear families (parents and children) and other small pedigrees have been used to estimate mutation and recombination rates in humans [1–4] and other species [5–7]. Pedigrees have informed breeding of domesticated animals [8], enabled the study of short-term evolution in natural populations [9], and can be used to study heritable diseases [10].

Genetic studies of rare, recessive traits pose a challenge to researchers when individuals expressing these traits are too sparse or too scattered to obtain sufficient genetic data. Endogamous populations with detailed pedigree records provide an important exception. Endogamous populations, defined by the practice of marrying within a social, ethnic, or geographic group, are often characterized by small effective population sizes with limited external admixture. These groups are of great interest to geneticists because a single small population can provide enough data to inform rare trait and rare variant studies with worldwide implications [11, 12]. Endogamous populations are also informative for common diseases [13, 14].

Extended pedigrees from endogamous populations provide a valuable system for studying heritable disease, but genetic data is typically limited to recent generations. If genetic information from every individual in the pedigree were available, we would be in a better position to understand the transmission of disease-associated variants throughout the history of the population. More specifically, we often know the disease phenotypes of ancestral individuals, but cannot obtain their genetic information. In these cases, reconstructed haplotypes would allow us to augment genome-wide association studies (GWAS), where large sample sizes are essential. In addition, reconstructed genomes would enable the computation of polygenic risk scores (PRS) [15, 16] for ancestral individuals.

Reconstructed ancestral haplotypes also allow us to study genome dynamics over short time scales, including inheritance patterns and haplotype transmission. In populations with large nuclear families, transmission distortion [17, 18] and other deviations from neutrality are particularly visible. Understanding which parts of the genome are over- or under-represented in the recent generations could help us identify forms of deleterious variation. From a theoretical perspective, there has been relatively little work on the question of how much ancestral reconstruction is possible given genetic information from contemporary individuals. One example from a small livestock pedigree can be found in [19].

Previous work on ancestral reconstruction has typically been applied to small pedigrees with no *loops* (marriage between close relatives). One of the earliest examples comes from the Lander-Green algorithm [20], which uses a hidden Markov model (HMM) with inheritance vectors as the hidden state and genotypes as the observed variables. Methods such as `SimWalk2` [21] and `Merlin` [22] use descent graphs and sparse gene flow trees (respectively) to extend the idea of likelihood-based computation to larger pedigrees. However, these methods do not perform reconstruction explicitly and also do not handle loops, as tree-based intermediate steps are common to both algorithms. With millions of loci and hundreds of individuals, the time complexities of these methods are prohibitive (see [23] for a runtime overview). Other HMM-based approaches such as `HAPPY` [24], `GAIN` [25], and `RABBIT` [26] reconstruct genome ancestry blocks, but do not tie them to specific individuals. `HAPLORE` [27] quantifies possible ancestral haplotype configurations but does not incorporate recombination, and the Bayesian approach in [28] is more suitable for haplotyping.

The authors of [29] reconstructed ancestral haplotypes for the purpose of identifying regions that contain susceptibility genes for schizophrenia. However, their pedigree was much smaller (with no loops), many fewer markers (450) were used, and several of the reconstruction steps were done by inspection or by hand, which does not scale to our scenario. Another study [30] reconstructed the African haplotype of an African-European individual who migrated to Iceland in 1802 and had 788 descendants, 182 of which were genotyped. However, this scenario is much simpler, as the regions of African ancestry within each descendant were easily identified and all belonged to the same individual.

The problem studied here is different from *pedigree reconstruction*, where genetic information is used to reconstruct (previously unknown) family relationships (see [31–36]). It is also different from ancestral reconstruction in a phylogenetic context, where a single tree represents the evolutionary relationships between species (see [37]).

In this study we apply our method to an Old Order Amish population from Lancaster, Pennsylvania who can trace their ancestry to founders who came from Europe to Philadelphia in the early 18th century (see Figure 3 of [38] for an analysis of the contributions of the 554 founders). The Amish are an ethno-religious group in the Anababtist tradition, with a history of detailed record keeping and marriage within the Amish community [39]. In this work, we study an unpublished pedigree of 1338 individuals, augmented [40] from a pedigree of 784 individuals originally described in the Amish Study of Major Affective Disorder [41, 42]. Roughly one third of the individuals in the original pedigree display some form of mood disorder, and about 19% have been diagnosed with bipolar disorder specifically [16]. Bipolar disorder in a broad sense is roughly 80% heritable in this pedigree [16], and recent work has focused on understanding the genetic basis of this disease [42]. The availability of genetic data from 394 contemporary individuals from this pedigree gives us an opportunity to use reconstruction as another lens on inheritance patterns of mood disorders.

Here we present a novel algorithm, `thread`, for reconstructing ancestral haplotypes given an arbitrary pedigree structure and genotyped or sequenced individuals from the recent generations. `thread` can be applied in a variety of scenarios including pedigrees with loops, inter-generational marriage, and remarriage. More ancestral chromosomes will be reconstructed as the percentage of individuals with genetic data increases, but our method can be applied even when this fraction is modest. This work represents a key step towards understanding the limits of quantifying the genomes of ancestral individuals in the absence of ancient DNA. `thread` is available as an open-source software package: https://github.com/mathiesonlab/thread.

## Materials and methods

### Problem statement

The first input to our reconstruction algorithm `thread` is a pedigree structure 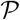. For each individual 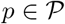 (aside from founders and married-in individuals), we have information about the mother *p*^(*m*)^ and father *p*^(*f*)^, which are also members of 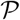. In the case of founders or married-in individuals, we represent *p*^(*m*)^ and *p*^(*f*)^ as 0’s. The pedigree may contain loops. Formally, a loop occurs when the undirected version of the directed marriage graph has a cycle (see [43, 44] for more information). Usually a loop means that the parents of a child share a common ancestor, but other loop structures are possible, e.g. when two brothers marry two sisters.

The second input is a dataset of phased haplotypes (e.g. in Variant Call Format, VCF) from a subset of individuals in the pedigree, typically from the most recent generations. Phasing assigns the alleles of each individual to parental haplotypes. Our aim is to reconstruct the haplotypes of as many ancestral individuals in the pedigree as possible. An illustration of the problem is shown in Fig 1.

**Fig 1.**
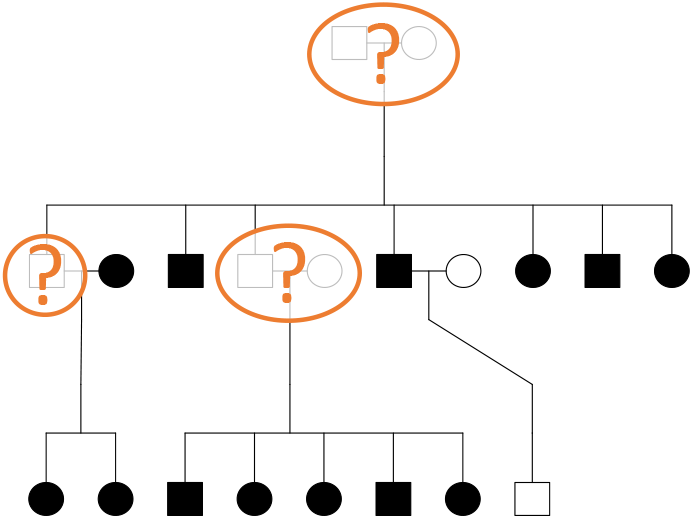
Problem statement illustration. Squares represent males and circles represent females. Horizontal lines create couples and show sibling relationships. Parents and offspring are connected by vertical lines. Filled in symbols represent individuals who have been genotyped. Our aim is to reconstruct all ungenotyped individuals (orange question marks) who have genotyped descendants.

### High level description

`thread` is built upon the idea of Identity-By-Descent (IBD). IBD segments are long stretches of DNA shared by a *cohort* of two or more individuals due to descent from a common ancestor (*source*). Our algorithm alternates between analyzing IBD segments and individuals. During each iteration (outer loop of Algorithm S1), we first consider each IBD segment independently (as opposed to working sequentially along the chromosome as an HMM would). We attempt to find the source of the IBD segment, as well as individuals who are on descendance paths from this ancestor to the cohort. After this step we proceed through each individual, clustering and assembling their associated IBD segments into haplotypes. During this grouping step we identify IBD segments that have been placed in a manner that is inconsistent with assigned haplotypes – in the next iteration we will update their common ancestors. We repeat these steps until no new haplotypes are reconstructed. Each chromosome of the genome is processed independently. A schematic of `thread` is shown in Fig 2, and pseudocode is given in Algorithm S1 (Supplementary Material).

**Fig 2.**
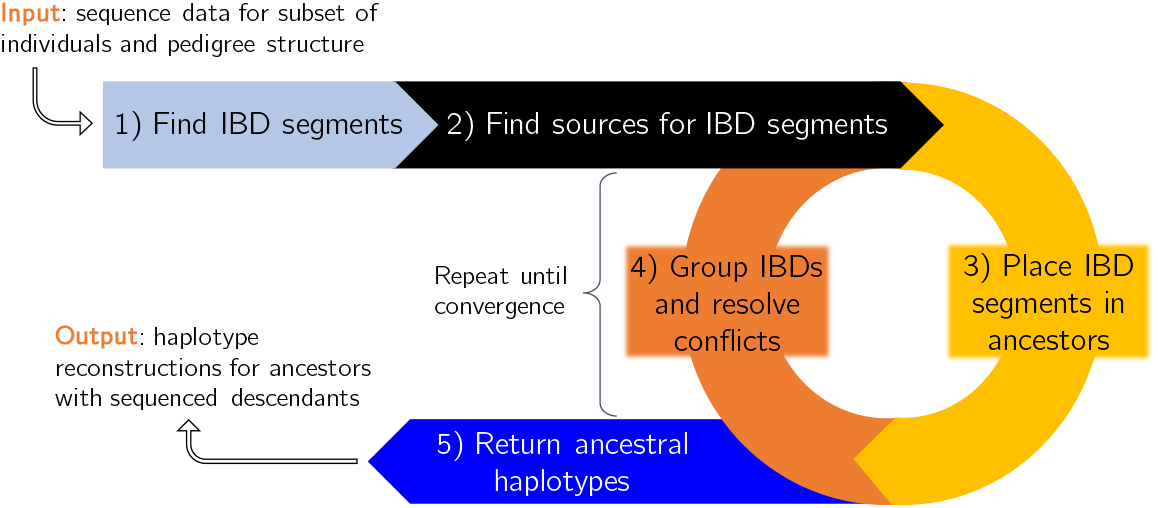
Algorithm overview. In the first two steps we identify IBD segments and find a list of potential sources for each one. In the iterative phase, we alternate between choosing sources for each IBD and grouping the IBDs that are placed within each individual. If the IBDs assigned to an individual can be arranged into two haplotypes meeting thresholds for coverage defined in Methods, then those haplotypes are considered ‘strong’. IBD segments that conflict with strong haplotypes are rejected and must be assigned a different source. When we are no longer building more haplotypes, we return the reconstructed chromosomes.

### Input pedigree

The Amish pedigree under study was developed from several sources, including the book *Descendants of Christian Fisher* [45], the Anabaptist Genealogy Database (AGDB) [40] and associated software `PedHunter` [38], and the Amish Study of Major Affective Disorder [41]. The AGDB is covered by an IRB-approved protocol at the NIH (protocol #97HG0192). All work contained within this study was approved by the IRB of the Perelman School of Medicine at the University of Pennsylvania (protocol #827037). A summary of the number of individuals in each generation is shown in Table 1, along with the number of genotyped individuals. The complete pedigree structure is shown in Figure S1 (created with the `kinship2 R` package [46]).

**Table 1.** Individuals per generation, with the founders assigned generation 1. The generation of a non-founder is defined to be one more than the maximum of the generations of their parents. Married-in individuals carry the generation of their spouse. The total number of individuals is 1338, with genotype information for 394 (largely in generations 9-12).

| Generation | # Individuals | # Genotyped |
| --- | --- | --- |
| 1 | 9 | 0 |
| 2 | 15 | 0 |
| 3 | 35 | 0 |
| 4 | 64 | 0 |
| 5 | 76 | 0 |
| 6 | 149 | 0 |
| 7 | 186 | 0 |
| 8 | 187 | 4 |
| 9 | 142 | 53 |
| 10 | 198 | 143 |
| 11 | 178 | 145 |
| 12 | 99 | 49 |

To assess the levels of relatedness and genetic similarity in this population, we compare the haplotypes of each pair of genotyped individuals. For each comparison, we compute the number of SNPs two haplotypes have in common over the total number of genotyped SNPs. Then for each pair of individuals, we take their overall genetic similarity to be the average of their haplotype similarities (paired to maximize similarity). We plot genetic similarity against kinship coefficient, which is computed using `PedHunter` and the AGDB (or our Amish pedigree if one or both of the individuals is not in the AGDB). The results are shown in Figure 3, which demonstrates that genetic similarity generally increases as kinship coefficient increases. We also include a histogram of inbreeding coefficients for each individual in Figure S2 (also computed using `PedHunter`). See [47] for more information about kinship and inbreeding coefficients.

**Fig 3.**
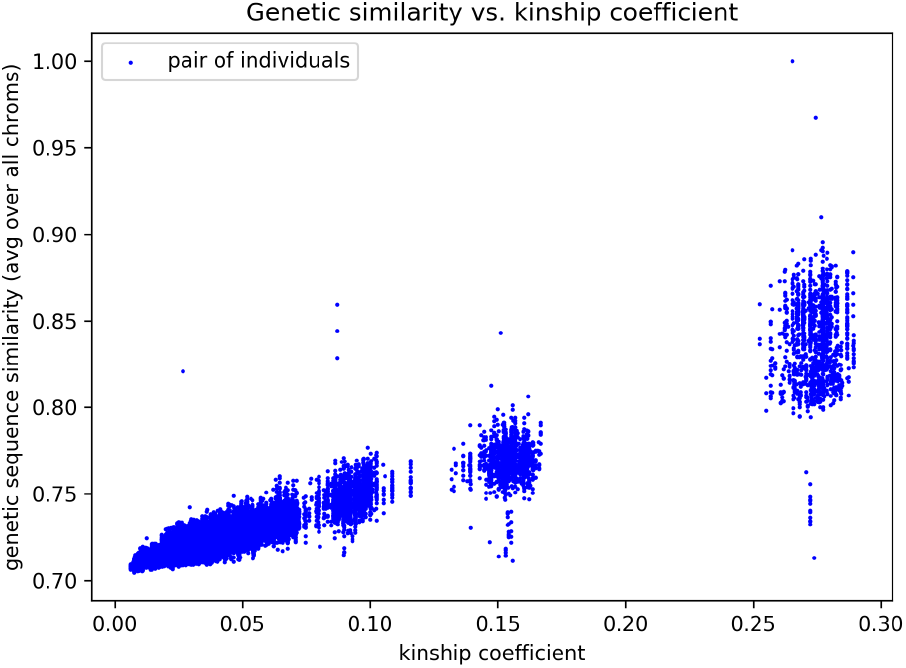
Genetic similarity vs. kinship. For each pair of genotyped individuals, we compute their genetic similarity (counting genotyped SNPs only) and plot this on the *y*-axis against their kinship coefficient on the *x*-axis. The expected linear trend is apparent, with an average sequence similarity of 72.5%. The minimum similarity out of all pairs was 70.5%, and the maximum was 99.999% (twins).

### Step 1: Find IBD segments

Step 1 begins by reading in the pedigree structure. The pedigree may contain inter-generational marriage and loops, and the individuals do not need to be separated into generations. Let *t* be the total number of individuals in the pedigree, *n* be the number of genotyped individuals, and *m* be the number of ungenotyped individuals with genotyped descendants. In the pedigree under study, *t* = 1338, *n* = 394, and *m* = 686, leaving 258 individuals with no genotyped descendants; we do not expect to be able to reconstruct these individuals.

Next, IBD segments between pairs of genotyped individuals are identified using `GERMLINE` [48], although `IBD-Groupon` for detecting IBD in groups could be used instead [49]. When using `GERMLINE`, we use the default parameters, except for the - haploid flag to indicate our dataset is phased. Thus the minimum IBD segment length is 3cM, and the length of seeds for exact matches is 128 markers. For each IBD segment *I*, we combine pairs until we obtain a cohort of individuals who share this segment, |*C*| ∈ {2, *n*}. The *descendance path* of an IBD segment includes all descendants of the source who also passed down the IBD to reach the cohort descendants. In this Amish pedigree, genotypes for each genotyped individual were obtained from Illumina Omni 2.5M SNP arrays, and then phased into haplotypes using `SHAPEIT2` [50]. The size of *C* ranged from two to 180 individuals. Table 4 shows the number of unique IBD segments found on each chromosome.

### Step 2: Find sources for IBD segments

In the next phase of `thread`, sources for each IBD segment are identified independently. By the end of this step, we will have enumerated all possible individuals who could have been the source of each IBD segment *I*, given its associated cohort *C*. This process is done only once and is not part of the iterative phase. When searching for *all* common ancestors of a cohort, each previous generation doubles the number of ancestors to search. `thread` maximizes efficiency in this exponential problem by merging overlapping paths using a modified breadth-first search algorithm (explained in detail below and in pseudocode in Algorithm S2).

First all the individuals in the cohort are added to a queue. For example, in Fig 4,

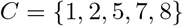

so we would start out with *Q* = (1, 2, 5, 7, 8). We then pop the first individual, *p*_0_ = 1 (in this example), off the queue. If *p*_0_ is an ancestor of all individuals in the cohort, we add *p*_0_ to a set of possible sources, *S*. Either way, we add *p*_0_’s parents to the back of the queue and keep processing individuals (even if *p*_0_ is an ancestor, its parents may be ancestors via paths that do not include *p*_0_). In this example, we first pop individual 1 off the queue. Since it has not been processed, we push 1’s parents onto the end of the queue to obtain

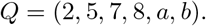

**Fig 4.**
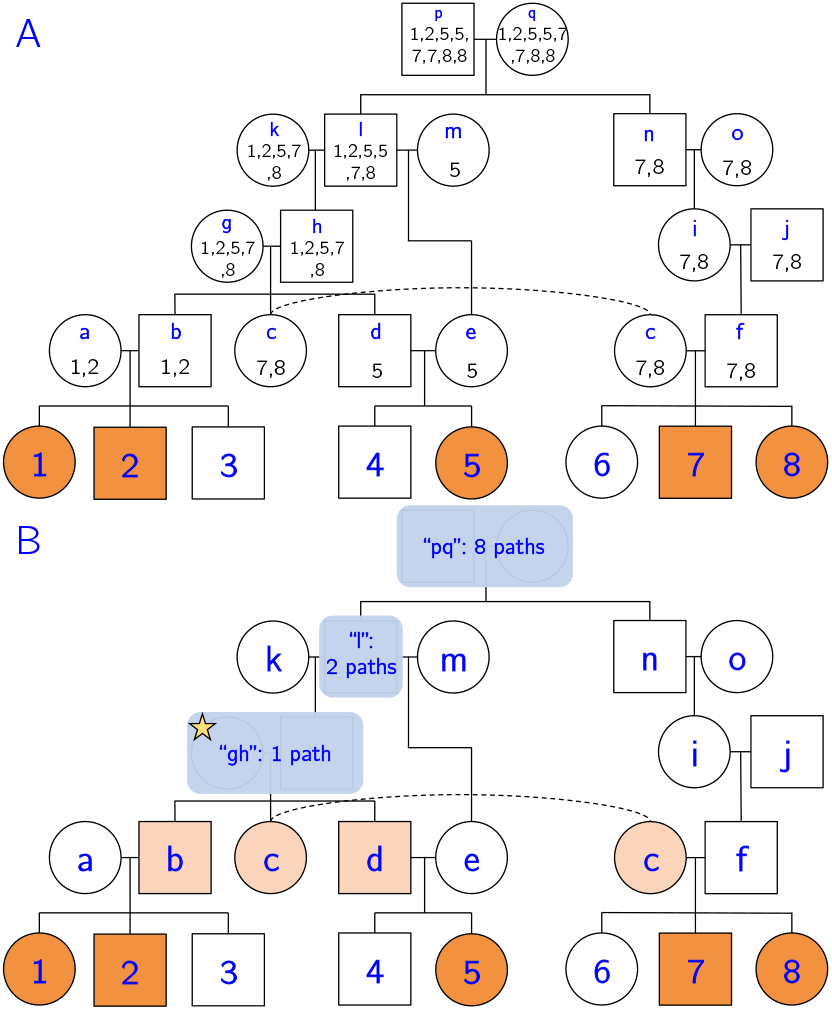
Source-finding illustration. A) Let individuals 1–8 be the genotyped individuals of this pedigree. Let *C* = {1, 2, 5, 7, 8} (orange individuals) be the cohort sharing IBD segment *I*. Note that this pedigree contains two loops, since *c* and *f* share recent ancestors *p* and *q*, and *d* and *e* share recent ancestor *ℓ*. The multiset *M_p_* for each ancestral individual *p* is shown below the node name. *M_p_* is formed by concatenating the multisets of *p*’s children, and it represents the number of paths from ancestor *p* to each member of the cohort. B) After trimming redundant ancestors and merging couples, we obtain a set of putative sources for the IBD segment. In this case, we have three potential sources: *S* = {*gh, ℓ, pq*}. We begin the iterative phase by selecting the source with the fewest descendance paths, which in this case is *gh* (starred). We place the IBD segment in individuals that are on all paths from *gh* to the cohort. In this case we would add the IBD segment to individuals *b*, *c*, and *d* (light orange).

Each time we add an individual *p* to the queue, we keep track of how many paths exist from *p* to the members of the cohort using a multiset *M*_*p*_. For the members of the cohort, *M*_*p*_ = {*p*} (just one path to themselves). When we add a parent to the queue, we concatenate the multisets of the individual’s children. For individual *a* in this example, its multiset would become *M*_*a*_ = {*1, 2*}, indicating one path to individual 1 and one path to individual 2. Going further up the pedigree, individual *ℓ* has two children, *h* and *e* with *M*_*h*_ = {1, 2, 5, 7, 8} and *M*_*e*_ = {5}. Concatenating these two multisets, we obtain the multiset *M*_*ℓ*_ = {1, 2, 5, 5, 7, 8}, indicating that there are two possible paths from *ℓ* to cohort member 5. As soon as an individual’s multiset contains all members of the cohort, the individual is a possible source.

There are two post-processing phases to the source-finding algorithm. (1) Trim redundant sources: a source is redundant if it is an ancestor of another source and does not add any unique descendance paths to the cohort. In other words, we do not want to include individuals if *all* their paths to the cohort go through another source.

Redundant sources do not depend on ordering effects, other than that children are added to the queue before their parents. Formally, if the cardinality of an individual’s multiset is equal to the maximum cardinality of the multisets of its children, it is redundant (for example, *k* is a redundant ancestor since |*M*_*k*_| = |*M*_*h*_|). (2) We merge couples into a single source, as typically we will not be able to resolve the source of an IBD segment beyond the couple level. Spouses with different multiset cardinality are an exception. These cases are usually caused by remarriage with at least one child from the second marriage. Individual *ℓ* is an example; we do not consider couple *kℓ* a source because |*M*_*k*_| < |*Mℓ*| due to *ℓ*’s remarriage to *m*. If the cardinalities had been the same (and not redundant), we would have considered *kℓ* a source.

In the Fig 4 example, we identify three potential sources: *S* = {*gh, ℓ, pq*}. Note that we cannot stop processing the queue when we get to source *gh*, as there exist sources further up the pedigree that are in previously unvisited descendance paths.

The use of multisets allows us to quickly determine the number of descendance paths from each source to the cohort. For each source *s* and each individual *c* in the cohort, let *m*_*s*_(*c*) be the multiplicity of *c* in *M_s_*. For example, in *M*_*ℓ*_, the multiplicity of individual 5 is two, meaning that there are two paths from *ℓ* to individual 5. The total number of descendance paths (*d*) from source *s* to cohort *C* (sharing IBD *I*) is the product of all the multiplicities:

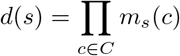

In this example, we obtain *d*(*gh*) = 1, *d*(*ℓ*) = 2, and *d*(*pq*) = 8. A few of these descendance paths are shown in blue in Fig 5 for clarity.

**Fig 5.**
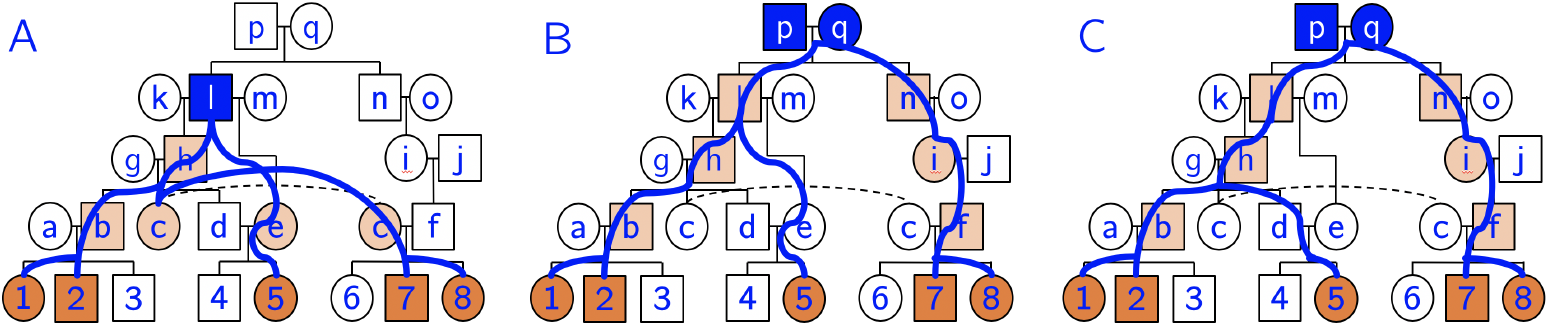
Example descendance paths. Given a cohort of five individuals sharing an IBD segment (orange), we often obtain multiple sources (blue nodes) and multiple descendance paths (blue lines) from each source. In this example we have 11 total paths from three sources. After we choose a source, we assign the IBD segment to ancestors along *all* descendance paths (light orange). A) One path from source *ℓ*. B-C) Two different descendance paths from the same source *pq*. We would not assign the IBD to *d* and *e* since they are not on all paths from this source.

Before moving into the iterative part of the algorithm, we take note of individuals that are on *all paths from all sources*. For example, individual *b* happens to be on all 11 paths from the sources, so we know that individual *b* should have the IBD segment.

### Step 3: Place IBD segments in ancestors

At this stage we begin the iterative part of the algorithm. Every iteration begins with lists of *reconstructed* individuals and *unreconstructed* individuals. During the first iteration, the *reconstructed* list only includes genotyped individuals. The goal of Step 3 is to select a source for each IBD segment out of the potential sources enumerated in Step 2. We use the greedy heuristic of choosing the source with the fewest paths, provided that it does not conflict with one of the *reconstructed* individuals. The intuition behind choosing the source with the fewest paths is that this source will (often) be more recent than others, with fewer meioses separating the source from the cohort. For example, in Fig 4, we would choose source *gh* since it has only one descendance path.

Once a source is selected, we can begin to look at the individuals that lie on paths from this source. In the case of only one path, all the individuals on the path will be given the IBD segment (*b*, *c*, and *d* in this example), thus augmenting the associated cohort. In the more common situation when we have multiple paths from the source, we give the IBD segment only to individuals that appear on *all* the paths. However, if we try to give this IBD segment to a *reconstructed* individual and it conflicts with both of the individual’s previously assigned haplotypes, we reject the source and choose the source with the next fewest paths. These tentative assignments result in potentially conflicting IBDs being assigned to the same individual, which we resolve in Step 4.

### Step 4: Group IBDs and resolve conflicts

During Step 3, we analyzed each IBD segment independently, identifying ancestral individuals who likely also share the IBD segment. In Step 4, we analyze the *individuals* independently and assemble the IBDs that have been placed within each individual. Say we are analyzing ancestral individual *p* with putative set of IBD segments 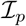. The goal of assembly is to separate the IBD segments into two haplotypes such that their sequences are consistent within each group. At a high level, this process can be compared to *de novo* genome assembly, where many small reads are stitched together to create contigs. We have an advantage over traditional assembly because we know the locations of each segment along the chromosome. But we may have included IBD segments that do not actually belong to a given individual, which we will need to identify and remove.

After grouping, we can identify individuals that are *reconstructed*. We define reconstructed as follows.

#### Definition 1.

*Reconstructed individual*. If we can group the IBDs placed within an individual into exactly two *strong* groups, we declare the individual *reconstructed*. Additionally, if there are two strong groups plus additional groups, the groups that are not strong must either have half as many IBD segments or be half as long as the weaker of the two strong groups.

#### Definition 2.

*Strong group*. To determine if a group is *strong*, it must meet a combination of thresholds: a minimum number of IBD segments and a minimum coverage (#SNPs reconstructed/#SNPs genotyped on the chromosome). We use a sliding scale: if the group contains 1-2 IBDs, it must cover 90% of the SNPs. If a group contains 3-9 IBDs, it must cover 70% of the SNPs. And if a group contains 10 or more IBDs, it must cover 50% of the SNPs. These parameters can be customized by the user.

Our grouping algorithm (covered in pseudocode in Algorithm S3) begins by identifying regions of homozygosity within the IBD segments. This is accomplished by condensing all segments in 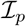 down into a single sequence with a list of alleles at each site. Any region greater than 300kbp with only one allele per site and at least 100 SNPs is declared homozygous. It is important to identify these regions early in the grouping algorithm, otherwise we may assume only one group contains this stretch. Each homozygous region is duplicated so that each chromosome will have a copy, and IBD segments contained within homozygous regions are not used in the next stages.

We process the remaining IBDs (those not incorporated into a group) one by one, from longest to shortest (in kbp). If the IBD does not overlap with any of the current groups, we create a new group initialized by the IBD segment. If the IBD does overlap with one or more groups, we add it to the group with the largest overlap (above a threshold).

At this point in the grouping algorithm, we have a set of homozygous groups, a set of heterozygous groups, and a set of remaining IBDs. If an IBD overlaps with two groups, we use it to merge the groups into one (assuming no previous overlap/conflict). Finally, we merge groups that “line up” with each other – i.e. they do not overlap, but their IBD segments span adjacent SNPs and were likely separated by an ancestral recombination event. At the end of this process, three situations may emerge:

- We have two clear groups (which meet our definition of *strong*) forming two haplotypes. This is the ideal scenario and it means we have a successful reconstruction of the individual.
- We have two strong groups, but we also have several weaker ones. This scenario is resolvable, as we can retain the two strong groups as the reconstruction, and reject the other groups. The IBD segments from the rejected groups give us a lot of information – since this individual was on *all* paths from the selected source, if the IBD segment does not fit with the reconstructed haplotypes, then we assume the source was incorrect. Throughout Step 4 we collect all IBD segments that have been incorrectly sourced to update in the next iteration.
- In all other situations, we typically cannot resolve the individual’s haplotypes. We may have only one group (which could be one of the individual’s haplotypes), but we do not declare the individual reconstructed. We could have many groups without two strong ones, or we may not have assigned any IBDs to the individual.

At the end of Step 4, we move individuals from the first two scenarios in to the *reconstructed* list. IBDs that did not cause any conflicts are marked as processed and we retain the rest to re-source in the next iteration. An illustration of the grouping algorithm is shown in Fig 6.

**Fig 6.**
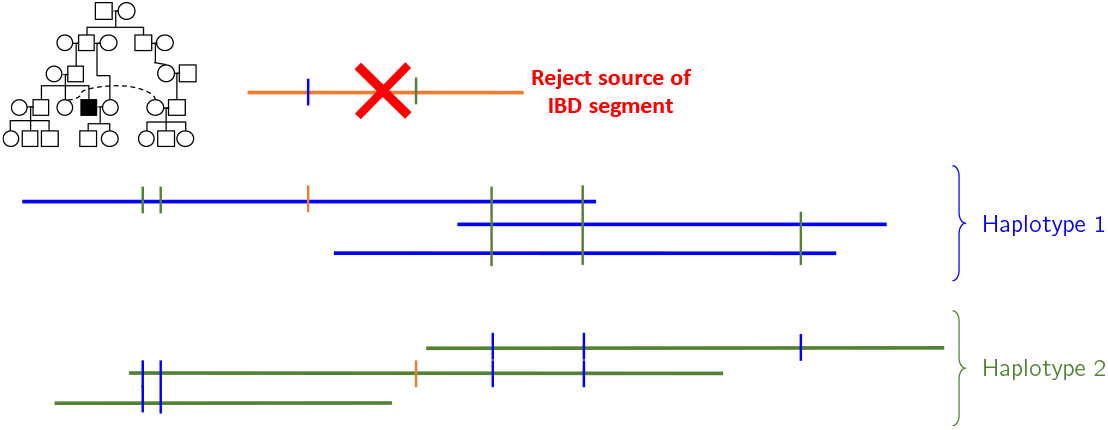
Grouping algorithm illustration. Each horizontal line represents one IBD segment that we placed within a specific individual (highlighted in the pedigree inset). Each vertical line indicates a difference (heterozygous site) between groups. In this case, the orange IBD segment conflicts with both the blue and green groups, so we would reject its source and attempt to find a new one in the next iteration.

### Step 5: Return ancestral haplotypes

At the end of Step 4 we have a set of IBD segments that were incorrectly sourced. We then repeat Step 3: we update the source for each such IBD by selecting the source with the next fewest paths. This allows us to assign the IBD to a new set of individuals. In the next Step 4 we treat reconstructed individuals and unreconstructed individuals differently. If an individual is already marked as reconstructed, we use each additional IBD to strengthen its groups or reject the new source of the IBD. If an individual has not been reconstructed, we run the grouping algorithm again. We keep iterating Steps 3 and 4 until we are no longer reconstructing new individuals.

The final step is to return the haplotype sequences for the reconstructed individuals. These may contain some gaps, but due to our coverage and length thresholds, if an individual is declared reconstructed, we will return at least half of each haplotype (for the chromosome under consideration). In the Supplementary Material we provide a theoretical complexity analysis of the full algorithm.

### Validation

We used two approaches to validate our approach directly. (1) To test the grouping algorithm, we grouped all IBD segments identified (by `GERMLINE`) as belonging to each *genotyped* individual. For this process, we did not specify which IBD segments belonged to which haplotypes, even though this information is known for genotyped individuals. (2) To evaluate our reconstruction results under the most realistic situation possible, we “left-out” one genotyped individual at a time and attempted to reconstruct that individual’s genome. We restricted this analysis to genotyped individuals with at least one genotyped descendant, which left 89/394 individuals. This approach can be interpreted as a form of leave-one-out cross-validation. In both of these validation approaches we compared the reconstructions to the original haplotypes using sequence similarity.

We also validated our method through whole-genome simulations. To mirror the demographic history of the Amish, we first simulate a founder population using msprime [51] with an effective population size of 10,000 diploids and the HapMap recombination map [52]. To simulate pre-migration endogamy in Europe, we create a 25-generation pedigree structure with a small population size (400 diploids). In this pedigree structure we constrain 5% of marriages to be between second-degree relatives (e.g. first-cousins), 10% between third-degree relatives, and 10% between fourth-degree relatives. We use `Ped-sim` [53] to simulate genetic data under this pedigree, using sequences from `msprime` as the founders. At the end of this pre-migration step, we are left with genetic data for 400 individuals, 186 of which we use as founders for the true pedigree structure after migration to North America. We again use `Ped-sim` to create genetic data for 1338 individuals in this pedigree, and retain this data for the same 394 individuals we have genotyped. Finally, we run `GERMLINE` and `thread` on this dataset, and compare the reconstructed sequences to their true values.

We were also interested in automatically predicting which individuals would be well reconstructed. To do this, we create a filter that identifies individuals based on their generation and average per-base IBD coverage, since we find that more recent individuals and those with high IBD coverage are typically more accurately reconstructed. In the Results section we highlight how many individuals pass this filter and the average reconstruction accuracy of these individuals.

This simulation framework provides the opportunity to test `thread` in a variety of scenarios. First, to model varying levels of endogamy, we varied the size of the pre-migration population from 200 to 2000 individuals. Additionally, we compared our original IBD source identification algorithm (greedy approach of taking the source with the fewest paths to the cohort) with a probabilistic approach (described in more detail in the Supplementary Material). In the probabilistic approach, we compute the probability that an IBD segment is transmitted from the source to all members of the cohort, taking into account the length of the IBD segment and the genetic map. We then take the source with the highest probability. If this source is rejected due to IBD conflicts in the grouping stage, we take the source with the next highest probability. Although we track the probability of IBD transmission throughout the paths from the source to the cohort, we do not account for the probability that non-cohort members do *not* have the IBD. Therefore these probability computations are not the probability of observing a particular configuration of the IBD segment, just the probability of transmission to the cohort.

By using IBD segments called by `GERMLINE`, the input to `thread` may already contain inaccuracies from segments that are Identical-By-State (IBS). To assess the impact of such inaccuracies, we used the true IBD segments as input to `thread`. These true IBD segments are identified by `Ped-sim` as part of simulating meiosis. These segments reflect IBD post-migration; there may be more ancient IBD sharing not reflected in the `Ped-sim` segments. In addition, it is possible that some individuals in the population share a known IBD segment due to IBS, but are not included in the `Ped-sim` segments because they did not inherit the segment from an ancestor in the pedigree.

Finally, this simulation framework allows us to experiment with some of the settings and thresholds in `thread`. We were particularly interested in the impact of the strong groups definition, as a single definition of strong group may not be applicable for individuals across the generations. To that end, we tried two different settings of the strong group definition.

- Setting A: For individuals with generation number 8 or greater, we use the same strong group settings as before. If the individual has generation number 7 or lower (i.e. more ancient) then we use the following sliding scale: if the group contains 1-3 IBDs, it must cover 90% of the SNPs. If a group contains 4-14 IBDs, it must cover 70% of the SNPs. And if a group contains 15 or more IBDs, it must cover 50% of the SNPs. This is meant to provide stricter thresholds for the more ancient generations.
- Setting B: For individuals with generation number 8 or greater, we use the same strong group settings as before. If the individual has generation number 7 or lower (i.e. more ancient) then we use the following sliding scale: if the group contains 1-3 IBDs, it must cover 70% of the SNPs. If a group contains 4-14 IBDs, it must cover 60% of the SNPs. And if a group contains 15 or more IBDs, it must cover 50% of the SNPs. This is meant to relax the IBD length thresholds, but still require more IBD segments for the more ancient generations.

## Results

### Grouping algorithm validation

After grouping the IBD segments called for each genotyped individual of the Amish pedigree, we analyzed the resulting haplotypes for coverage and correctness. Fig 7 shows two chromosomes of a *genotyped* individual that were reconstructed using the grouping algorithm `thread`. Each horizontal line represents one IBD segment shared with a cohort of other genotyped individuals. IBD segments of the same color represent haplotypes, and have a consistent sequence along the chromosome. In other words, if we agglomerated the IBD segments of a single color in Fig 7A or 7B, a single sequence would emerge. In general we found that our grouping algorithm worked very well for genotyped individuals, who typically share many IBD segments with other members of the pedigree. In 8558 out of 8668 chromosomes (22 per genotyped individual), we successfully grouped placed IBDs into two haplotypes. Very occasionally (1.3% of cases) we obtained three groups (example in Fig 7A). In chromosome 21 for one individual, all IBD segments belonged to a single group, indicating homozygosity along the entire chromosome. Looking at the sequences of these grouped haplotypes, they are very close to the true haplotypes. Average per chromosome accuracies are between 98.6% and 99.7%, as measured by sequence similarity.

**Fig 7.**
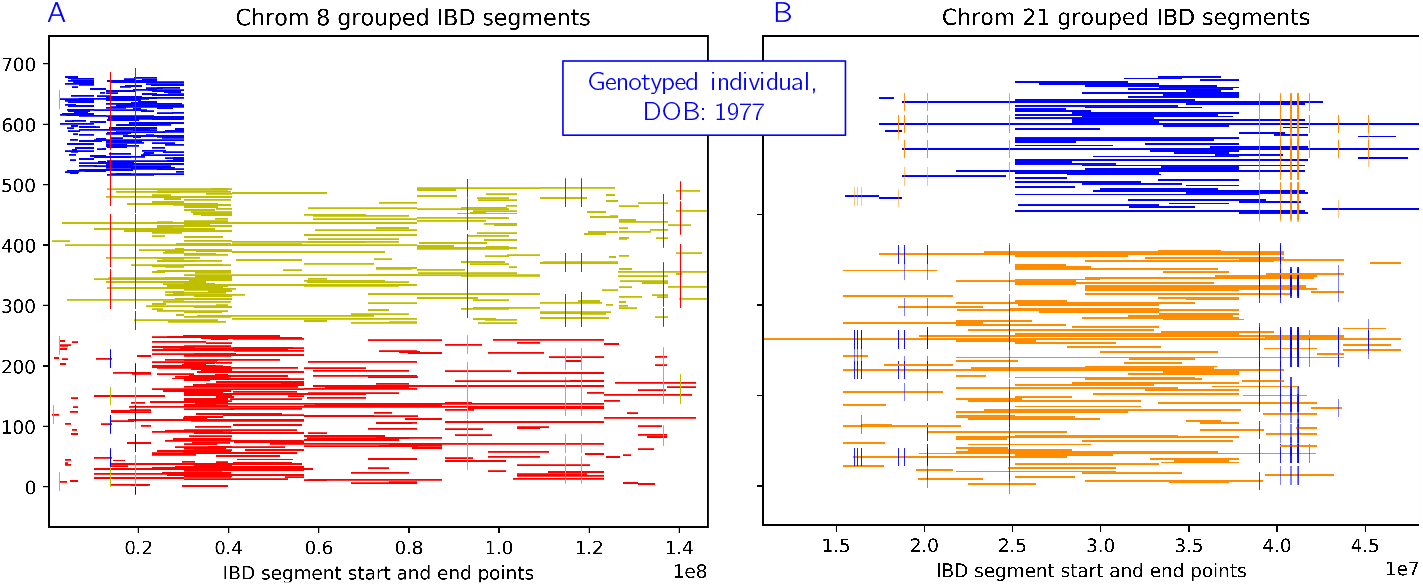
Example of the grouping algorithm on a genotyped individual. Each horizontal line represents one IBD segment shared with a cohort of other genotyped individuals. IBD segments of the same color represent haplotypes, and have a consistent sequence along the chromosome. Small vertical lines represent heterozygous sites between the two haplotypes. A) Chrom 8: very occasionally we merge groups incorrectly and obtain three groups. B) Chrom 21: we almost always see two clear haplotypes (here we also see a large stretch of homozygosity).

### Leave-one-out validation

To evaluate our reconstruction results under a realistic situation, we “left-out” one genotyped individual at a time and attempted to reconstruct that individual’s genome. We restricted this analysis to genotyped individuals with at least one genotyped descendant, which left 89/394 individuals. On average, these individuals had 9.87 genotyped descendants (min 1, max 48). Table 2 shows the results of this procedure for chromosomes 18-22. For example, in the case of chromosome 18 we reconstructed 71/89 individuals, with an average sequence similarity of 96.7% (measured against the corresponding true sequences). To be *reconstructed*, at least half the chromosome must be assembled for each haplotype, with sufficient support in terms of coverage and number of IBD segments (further details in **Definitions 1–2**). It is likely that we reconstructed parts of the remaining 18 individuals, but they did not meet our definition of reconstructed.

**Table 2.**
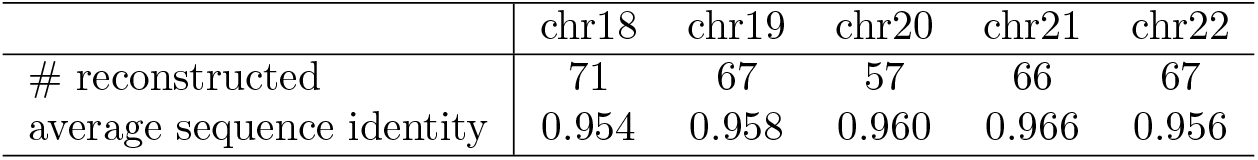
Leave-one-out results. For each chromosome 18-22, we left out one genotyped individual in turn and attempted to reconstruct their haplotypes. The second row shows how many individuals (out of 89) met our criteria for reconstructed. The third row shows the average sequence identify of the individuals we we were able to reconstruct, measured against their true sequences.

|  | chr18 | chr19 | chr20 | chr21 | chr22 |
| --- | --- | --- | --- | --- | --- |
| # reconstructed | 71 | 67 | 57 | 66 | 67 |
| average sequence identity | 0.954 | 0.958 | 0.960 | 0.966 | 0.956 |

We also explored the relationship between number of genotyped descendants and the probability of being reconstructed. We computed *P* (reconstructed | *D*_*g*_), where *D*_*g*_ was *low*, *medium*, and *high* based on breaks in the distribution of number of genotyped descendants. We did not find a strong relationship, perhaps because the chance of being reconstructed depends more on the pedigree structure itself and the likelihood that a particular individual can be identified as the source of an IBD segment. To address this hypothesis, we computed the Jaccard similarity coefficients between each pair of chromosomes (from 18-22). For this analysis we considered the data for each chromosome as a binary vector of length 89, with each individual being reconstructed or not. Overall the Jaccard coefficients were high (all pairs between 0.66 and 0.87, with an average of 0.79), indicating that an individual’s position in the pedigree structure is an important factor when considering our ability to reconstruct their genome.

### Simulation results

After running `thread` on the entire genome for the simulated data described above, we assess the accuracy by comparing the reconstructed genomes to the correct simulated ones. The average accuracy of the reconstructions is shown in Table 3. We note that although we simulated a small population with marriage between close relatives, the number of unique IBD segments per chromosome is less than half the observed number of unique IBD segments in the genotyped Amish individuals, which likely led to fewer reconstructed individuals.

**Table 3.** Whole-genome ancestral reconstruction results: simulated data. The second column shows the total number of IBD pairs identified between genotyped individuals. The third column shows the number of unique IBD segments per chromosome. The fourth column shows how many iterations the algorithm needed to converge. The fifth column shows the number of ancestral (ungenotyped) individuals we were able to successfully reconstruct. The sixth column shows the average sequence similarity of the individuals we were able to reconstruct, as compared to their true genomes. The rightmost two columns show the number of individuals that we *predicted* would be very well reconstructed, along with their average accuracies.

| chr | IBD pairs | unique IBDs | iter | recon. | acc. % | filter | filter acc. % |
| --- | --- | --- | --- | --- | --- | --- | --- |
| 1 | 312526 | 10447 | 8 | 181 | 86.05 | 34 | 90.19 |
| 2 | 314518 | 9588 | 9 | 173 | 86.14 | 40 | 89.42 |
| 3 | 273032 | 8006 | 6 | 162 | 86.52 | 33 | 91.39 |
| 4 | 186735 | 7336 | 5 | 164 | 86.42 | 42 | 89.95 |
| 5 | 258649 | 7891 | 9 | 195 | 86.22 | 30 | 90.25 |
| 6 | 225494 | 7249 | 8 | 189 | 85.97 | 45 | 89.94 |
| 7 | 198285 | 6657 | 6 | 172 | 85.81 | 37 | 89.97 |
| 8 | 169540 | 5833 | 8 | 143 | 86.64 | 29 | 92.55 |
| 9 | 188457 | 5687 | 5 | 127 | 85.31 | 28 | 91.15 |
| 10 | 209888 | 6191 | 6 | 146 | 87.97 | 33 | 92.16 |
| 11 | 183196 | 5910 | 5 | 163 | 87.25 | 31 | 91.50 |
| 12 | 212065 | 6040 | 5 | 168 | 86.22 | 37 | 89.12 |
| 13 | 136549 | 4561 | 7 | 156 | 86.62 | 28 | 91.19 |
| 14 | 121796 | 3858 | 6 | 132 | 86.63 | 29 | 91.96 |
| 15 | 157815 | 4837 | 5 | 169 | 87.52 | 27 | 92.86 |
| 16 | 150385 | 4515 | 6 | 134 | 88.34 | 29 | 92.70 |
| 17 | 136973 | 4351 | 6 | 148 | 87.34 | 30 | 90.65 |
| 18 | 112425 | 3885 | 7 | 158 | 86.74 | 28 | 90.54 |
| 19 | 120179 | 3523 | 4 | 125 | 86.82 | 23 | 88.61 |
| 20 | 135525 | 3959 | 5 | 143 | 87.40 | 23 | 90.52 |
| 21 | 79550 | 2109 | 6 | 90 | 88.73 | 17 | 92.96 |
| 22 | 86547 | 2528 | 4 | 85 | 86.90 | 17 | 90.27 |

Individual level results are shown in Figure 8. Here we plot two reconstruction metrics that increase as reconstruction is more successful – the number of chromosomes reconstructed and the average accuracy of the reconstructed chromosomes. The strongest relationship appears with generation, where more recent individuals generally have more complete and accurate genome reconstructions. There is a weak relationship with the number of genotyped direct descendants (children and grandchildren). Individuals with more genotyped direct descendants generally have more accurate reconstructions, but not necessarily more chromosomes reconstructed. As inbreeding coefficient increases we see a general increase in accuracy, but again not a strong relationship. We also show the relationship between generation, reconstruction accuracy, and number of chromosomes reconstructed in a more holistic fashion in Figure S3. In addition, we demonstrate a few examples of how IBD segments are removed during conflict resolution in Figure S4. In the first example we show how incorrectly placed IBD segments were successfully identified and removed, leading to a perfect reconstruction. In the second case we show an example where `thread` failed to identify incorrect segments, leading to a reduced reconstruction accuracy.

**Fig 8.**
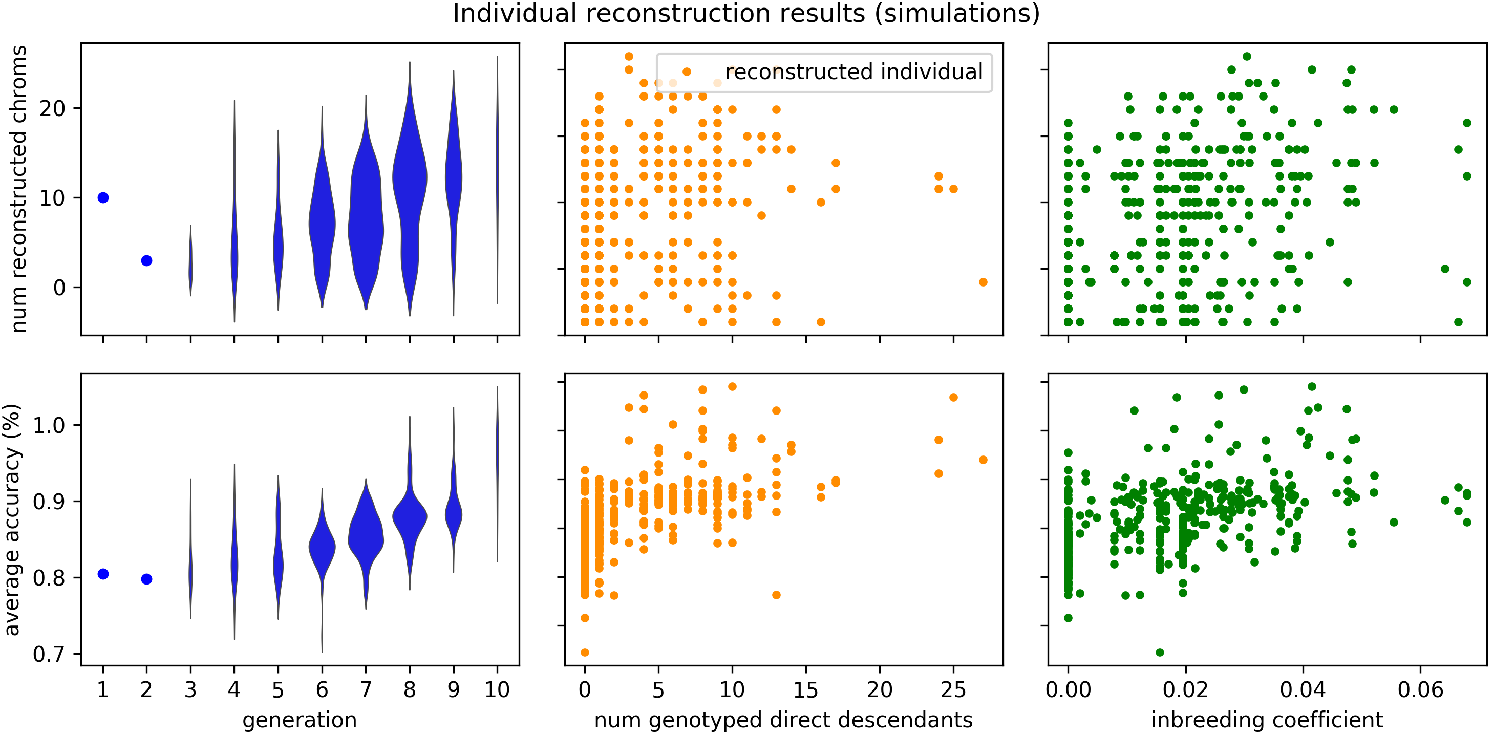
Individual results: simulated data. The same results as Table 3, but shown on the individual level. The top set of figures shows reconstruction completeness as measured by the number of reconstructed chromosomes. The bottom set of figures shows reconstruction accuracy as measured by sequence identity averaged over the reconstructed chromosomes. These two metrics are plotted against three statistics about each individual: the generation number (lower is more ancient), the number of genotyped direct descendants (children and grandchildren), and the inbreeding coefficient as calculated by `PedHunter` using the entire AGDB comprised of more than 500,000 individuals.

The results of varying pre-migration population size and IBD source selection algorithms for chromosome 21 are shown in Figure 9. The greedy algorithm is denoted “min path” and the probabilistic algorithm is denoted “max prob”. As the population size increases, we generally see more accurate reconstructions, but fewer individuals reconstructed. This tradeoff is likely due in part to differences in cohort size – if only a few individuals share an IBD segment, its source can be more easily identified than if many individuals share the IBD. This leads to more accurate reconstructions, but fewer individuals will be assigned sufficient IBD segments. The probabilistic source identification algorithm typically results in fewer reconstructed individuals, but higher accuracies (up to 98.38%). By definition the “max prob” source will have at least as many paths as the “min path” source, but there will be fewer individuals on all paths from the source to the cohort. This leads to fewer IBD segments placed into individuals, but higher accuracy for those that are placed.

**Fig 9.**
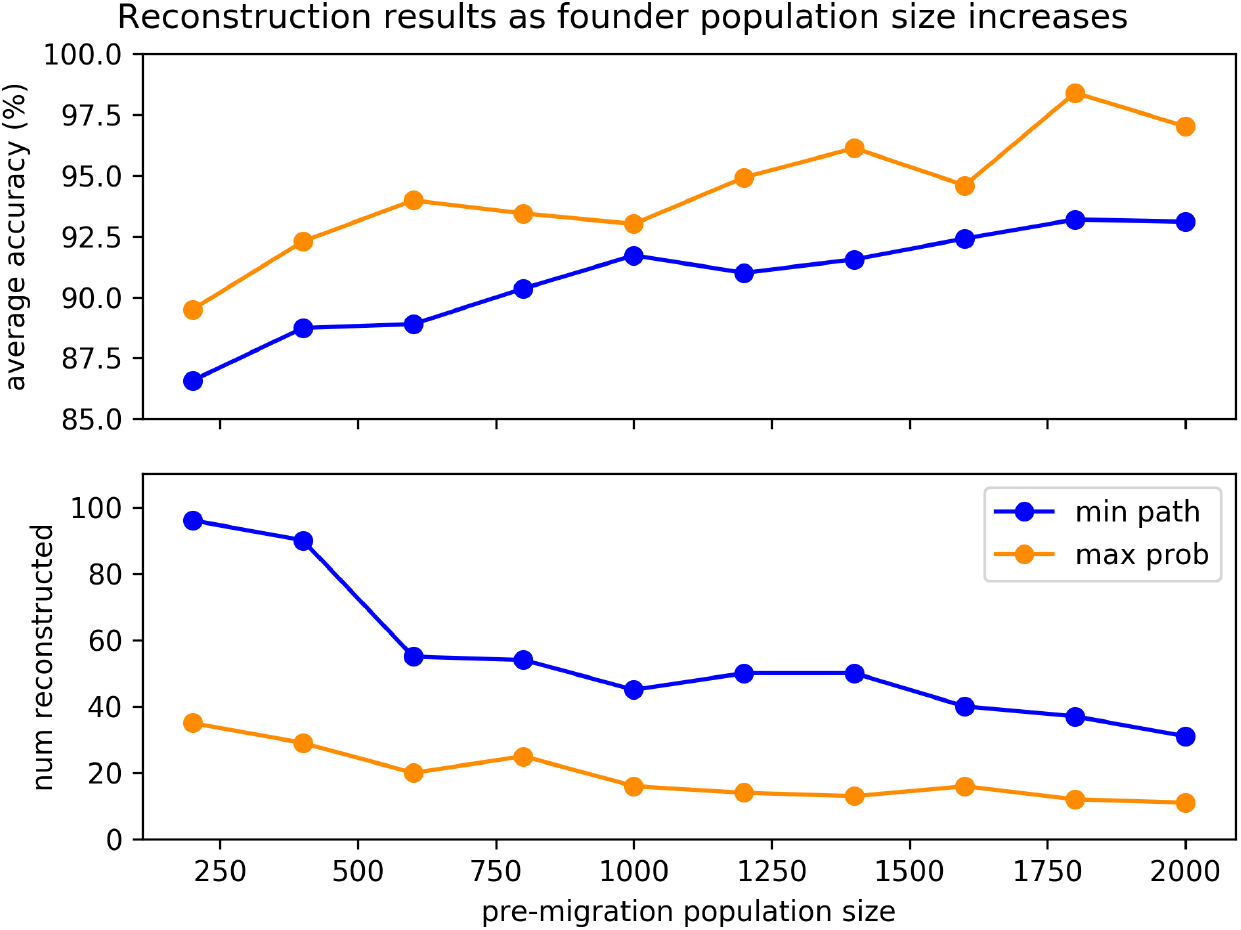
Varying population size and source-finding approach: simulated data. The top panel shows the average reconstruction accuracy of chromosome 21 as a function of pre-migration population size. The bottom panel shows the number of reconstructed individuals for the corresponding scenarios. The greedy source-identification algorithm is denoted “min path” and the probabilistic algorithm is denoted “max prob”. There is a clear tradeoff between accuracy and the number of individuals reconstructed.

The results of using the true IBD segments from `Ped-sim` are shown in Table S1, for chromosomes 18-22. Overall there were more unique IBD segments than called by `GERMLINE`, possibly due to more accurate detection of the endpoints of IBD segments. Overall however, the results do not show dramatic improvement, and the number of reconstructed individuals is typically fewer than when using `GERMLINE` (but not less accurate). This indicates that IBS called as IBD is not likely a major cause of inaccuracies in `thread` reconstructions. (For IBS segments with large cohort sizes, if no sources can be identified then we skip the segment.) Based on the results in Figure 9, source confusion likely plays a bigger role in creating reconstruction error.

Finally, the results of modifying the strong group criteria are shown in Table S1, for chromosomes 18-22. Setting A reduces the number of reconstructed individuals, which makes sense since it is a strictly stronger criteria than before. But gains in accuracy are minimal. Setting B produces similar results as the original settings, indicating that much more relaxed criteria are needed to reconstruct individuals in more ancient generations (which would likely lead to decreases in reconstruction completeness and accuracy). Overall, ancient individuals remain difficult to reconstruct.

### Reconstruction results

After running `thread` on each chromosome using the entire Amish pedigree and all genotyped individuals, we assessed the results in terms of how many individuals were successfully reconstructed. For all chromosomes, `thread` converged in 5-10 iterations, and the number of successfully reconstructed ancestral individuals ranged between 162 and 248 (24%-36% of the 686 individuals with genotyped descendants). On average this is 224 individuals per chromosome, with an overall average of 79% of their haplotypes reconstructed. See Table 4 for the details of each chromosome. Using a single CPU, the runtime was always less than 24 hours for a single chromosome. Typical memory requirements were 40G for all but the largest few chromosomes, which needed 80G.

**Table 4.** Whole-genome ancestral reconstruction results: Amish data. The second column shows the total number of IBD pairs identified between genotyped individuals. The third column shows the number of unique IBD segments per chromosome. The fourth column shows how many iterations the algorithm needed to converge. The fifth column shows the number of ancestral (ungenotyped) individuals we were able to successfully reconstruct. We require a successfully reconstructed chromosome to have two haplotypes that cover at least half the chromosome, with sufficient IBD support for each haplotype. Finally, the last column shows the runtime.

| chr | IBD pairs | unique IBDs | iter | reconstructed | time (hrs) |
| --- | --- | --- | --- | --- | --- |
| 1 | 351230 | 28359 | 10 | 248 | 22.91 |
| 2 | 288059 | 26962 | 7 | 248 | 19.19 |
| 3 | 246909 | 22488 | 6 | 232 | 11.23 |
| 4 | 250878 | 20980 | 6 | 223 | 9.83 |
| 5 | 219746 | 19448 | 7 | 236 | 9.14 |
| 6 | 244751 | 20883 | 7 | 224 | 10.76 |
| 7 | 225155 | 19370 | 6 | 216 | 6.73 |
| 8 | 222422 | 16950 | 6 | 241 | 4.96 |
| 9 | 232309 | 17547 | 6 | 236 | 3.85 |
| 10 | 220391 | 16822 | 7 | 223 | 6.22 |
| 11 | 198145 | 15416 | 8 | 238 | 5.49 |
| 12 | 175552 | 16712 | 7 | 227 | 5.08 |
| 13 | 161721 | 13296 | 5 | 202 | 2.74 |
| 14 | 169906 | 11867 | 6 | 239 | 2.36 |
| 15 | 180198 | 12179 | 7 | 248 | 2.60 |
| 16 | 143985 | 13010 | 7 | 251 | 2.74 |
| 17 | 121688 | 11768 | 5 | 195 | 2.44 |
| 18 | 129719 | 11359 | 7 | 219 | 2.13 |
| 19 | 141857 | 10702 | 6 | 218 | 1.44 |
| 20 | 139179 | 9910 | 8 | 235 | 1.95 |
| 21 | 77745 | 5020 | 6 | 175 | 0.68 |
| 22 | 73766 | 5773 | 5 | 162 | 0.77 |

We also compare the IBD length distributions of the simulated data and the real data, as shown in Figure 10. In general the simulations have fewer unique IBDs (especially short IBDs), but the distribution shapes are similar.

**Fig 10.**
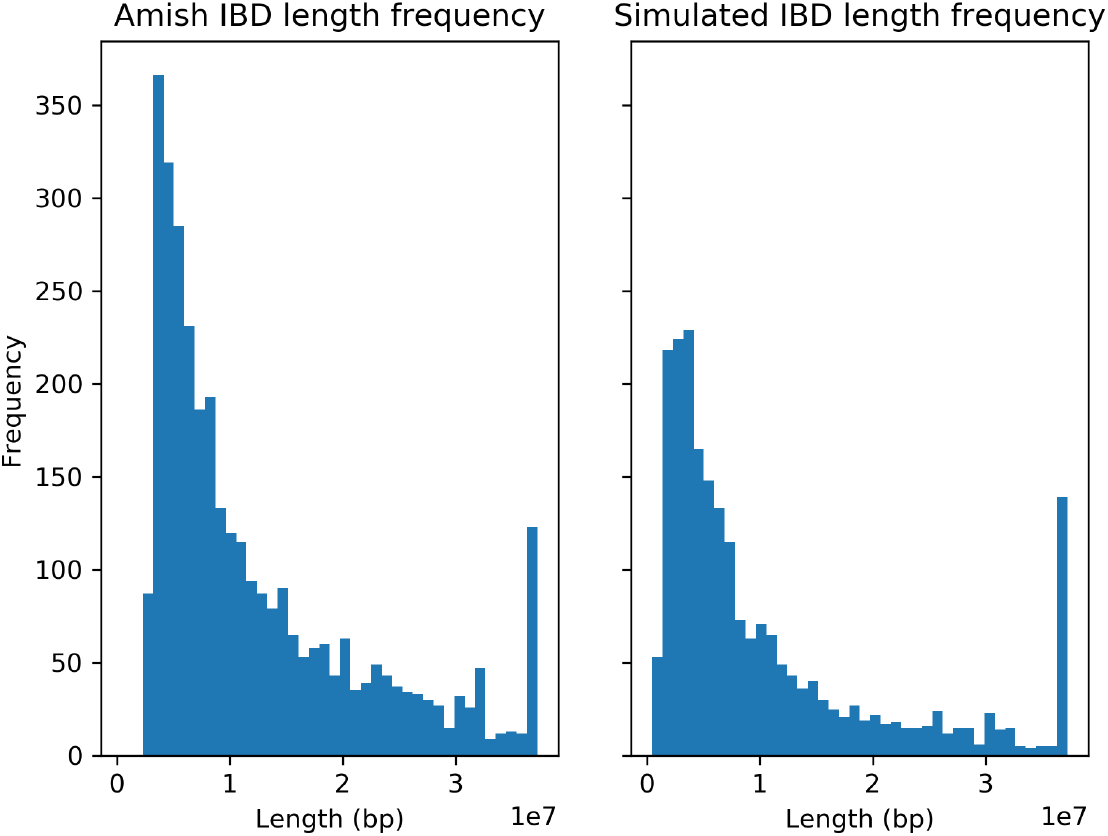
IBD length distribution. Left: IBD length distribution for the real data for chromosome 21. Right: IBD length distribution for the simulated data for chromosome 21. *x*-axis units are 10Mbp.

The conflict resolution step was essential for removing misplaced IBD segments and routing them to other sources. An example is shown in Figure 11. In this case, the green and blue groups were removed from this individual, as they were much less *strong* than the cyan and red groups. In the next iteration, we re-source the associated IBDs and consider the individual reconstructed. Examples of successful ancestral reconstructions are shown in Figure 12, for a variety of different chromosomes and generations back in time. As expected, in the more distant generations, we place fewer IBD segments and generally have less coverage over the chromosome.

**Fig 11.**
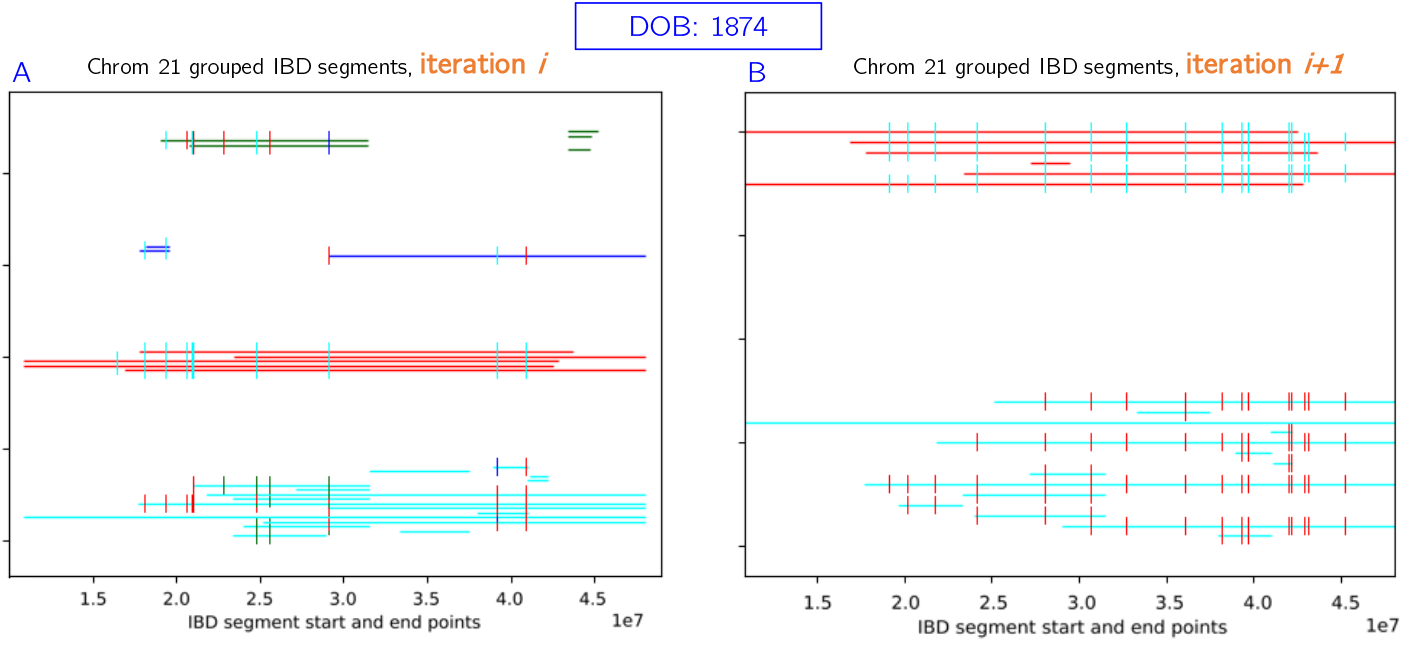
Conflict resolution example. The blue and green groups are removed, since they are less *strong* than the cyan and red groups. In the next iteration, we retain only strong groups and consider the individual reconstructed. Newly sourced IBDs after this point may not conflict with these reconstructed haplotypes.

**Fig 12.**
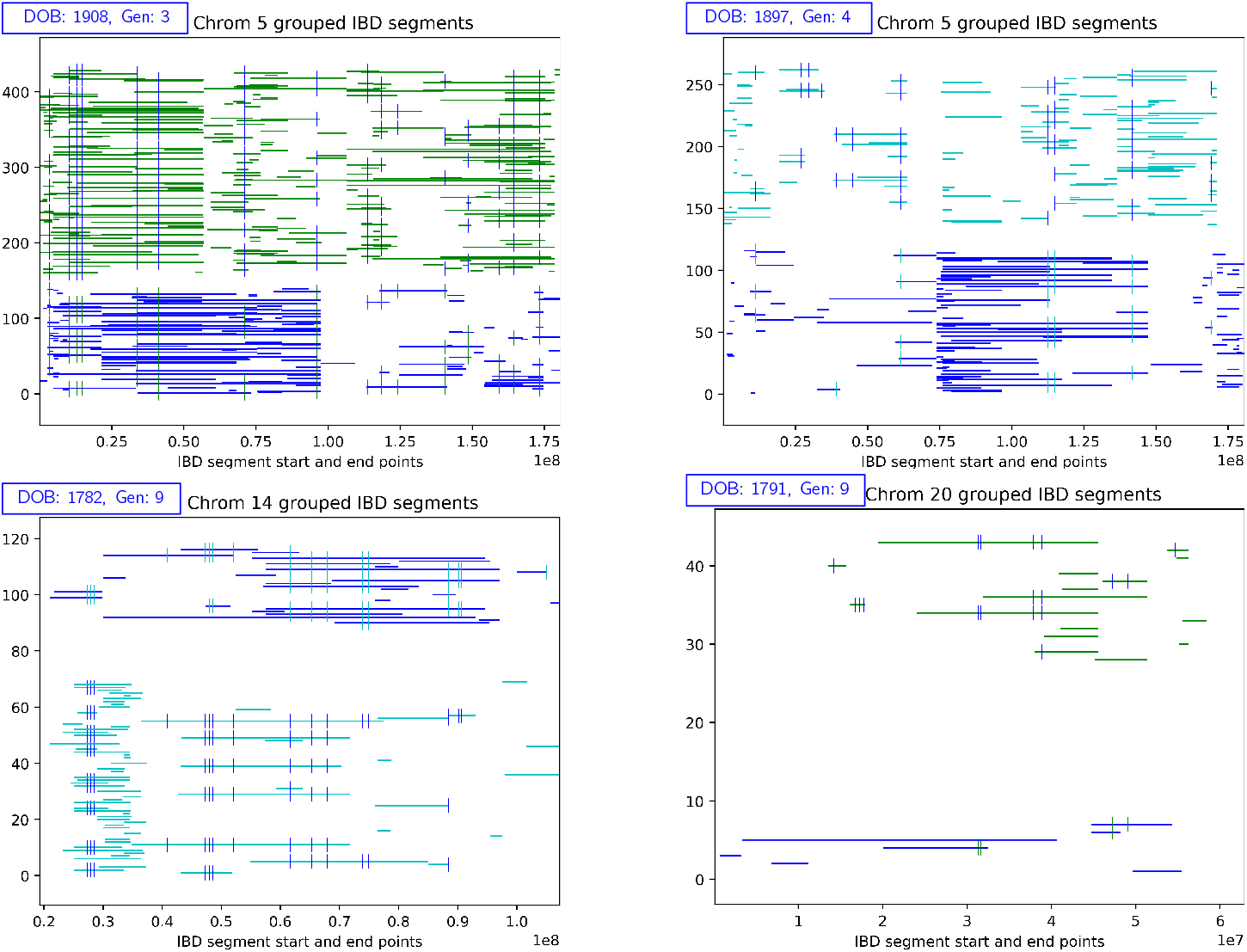
Successful ancestral reconstructions. Ancestral reconstructions of ungenotyped individuals, from a variety of chromosomes and generations (back in time). As we go back in time, we generally have fewer IBD segments to group.

Although we reconstruct many individuals well in the recent generations, there are many haplotypes we are unable to resolve. Several examples are shown in Figure S6. Sometimes `thread` constructs one haplotype successfully, but not the other (Figure S6A). Often we have some successful reconstruction, but the groups do not meet our threshold for “two strong” since the third group has too many IBD segments (Figure S6B). Sometimes there are four groups, which could represent ambiguity between the individual’s spouse or close relative (Figure S6C). Sometimes there are too many IBD segments placed within an individual, which could arise if they have many descendants (Figure S6D).

Table 4 and Figure 13 show our results in a holistic view. Table 4 shows how many individuals we are successfully reconstructing for each chromosome. Figure 13 shows these same results on the family level, broadly indicating which individuals we are reconstructing well. Figure S5 shows these results on the individual level.

**Fig 13.**
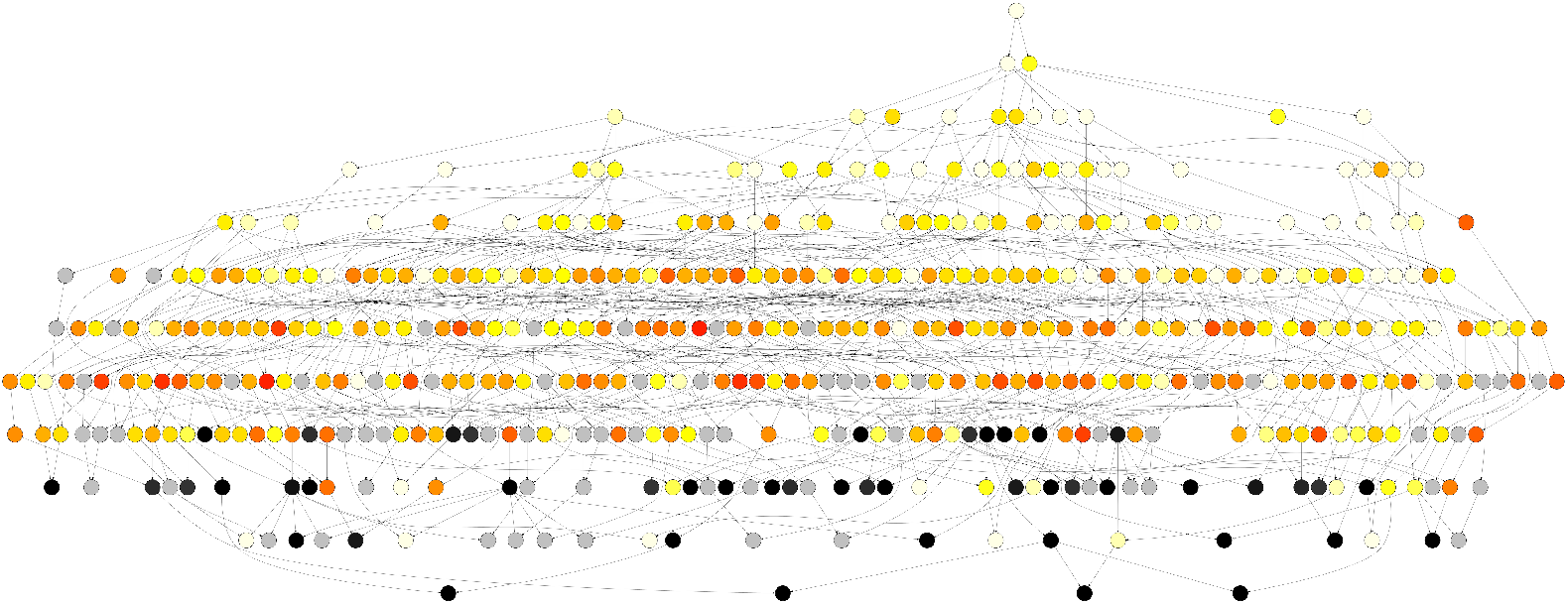
Nuclear family graph. Each node represents a nuclear family (parents and children). When a child of one family becomes the parent of another, we draw an edge. Black nodes have at least 80% of the family genotyped. Gray nodes have at least 80% of the family without genotyped descendants. Yellow (fewer) – Red (more) colors represent the average number of chromosomes reconstructed for the individuals in the family.

### Lost regions

We were particularly interested in regions of the genome that have been “lost” over time, as they may be deleterious. To assess this, we looked for regions of the genome with a low fraction of reconstructed individuals. We measured this by analyzing non-overlapping sliding windows of 50 SNPs; if less than 40% of the bases in a window were not reconstructed, we declared the region difficult to reconstruct. We then merged adjacent regions to form a set of “lost” regions. The telomeres of each chromosome formed the majority of regions that were difficult to reconstruct, which is expected. Across the entire genome, there was only one non-telomeric region that met our criteria for a lost region, chr16:70824983-71066078. This region on chromosome band 16q22, containing most of the large gene *HYDIN* (chr16:70835987-71264625), is known be genomically unstable in two ways that could make it difficult to reconstruct haplotypes. The more likely difficulty is that there are frequent segmental duplications with varying breakpoints in this region [54], which have been associated with autism [55].

Re-examination of the copy-number variants (CNVs) called in a previous study [56] by RLK using the software `PennCNV` [57] shows that there are overlapping intervals of a duplication within the interval chr16:70696689-70745016 in 67 genotyped individuals in the pedigree. One boundary of the CNV is shared by most of the 67 individuals. The other boundary, however, varies considerably, which is unusual for an inherited CNV and may reflect either the difficulty in calling genotypes mentioned above or genomic instability. The average length of the duplication among the 67 individuals is 32,641 bp.

The less likely difficulty is that in evolution, the instability of this region led to a recent (by evolutionary time scales) paralogous duplication of the region that landed on human chromosome band 1q21, and contains the paralogous gene *HYDIN2*. The chromosome 1 copy is sufficiently similar to the 16q22 that it caused difficulty in assembling early versions of the human genome [58]. The sequence similarity of the 16q22 and 1q21 segments could lead to imperfect genotypes in either interval.

## Discussion

The methodology behind `thread` represents a new direction for ancestral reconstruction that scales in both the number of individuals and the number of loci. Previous ancestral haplotype reconstruction algorithms have either been too slow to apply, too rigid to accommodate a complex pedigree, perform steps by hand, or consider a more diverse ancestral population. Although a likelihood approach to reconstruction is theoretically possible, our work represents a practical alternative as pedigree size and complexity continues to grow. We note that our method is most suitable when genotyped individuals exhibit high levels of IBD sharing, ideally with each IBD segment descending from a single ancestor. `thread` may need further development to handle pedigrees from populations with very large effective population sizes and/or high levels of admixture.

Through simulations we validate `thread` in a variety of scenarios, including a range of ancestral population sizes. With realistic simulation parameters it was difficult to obtain the number of unique IBDs found in the Amish population, which is likely a reason why we reconstruct fewer individuals in simulations. One overall trend is that as the population size increases, we reconstruct fewer individuals, but their reconstructions are more accurate. Well reconstructed individuals also tend to be closer to the present. A future version of the algorithm could use these observations to build up reconstructions gradually – after an initial group of individuals is accurately reconstructed, they could be added to the original list of “genotyped” individuals, then the entire algorithm could be run again to reconstruct a next group of individuals, and so on. This type of approach might be particularly useful for populations with less endogamy, where reconstructing ancient individuals is naturally more difficult.

There are many possible algorithmic improvements to the IBD-based approach of `thread`. The grouping algorithm could make use of the genetic map to merge groups at recombination hotspots. More realistic simulations could model crossover interference and sex-specific recombination maps, as in [53]. In rare cases (only chromosome 21 for one genotyped individual) the entire chromosome is homozygous, so the criteria of two strong groups could be relaxed or made more flexible. In terms of implementation, `thread` could be parallelized across IBD segments and individuals.

An important use case for genealogies of isolated populations has been to study rare recessive diseases, including finding the causative genes. After collecting data on living affected individuals, it may be of interest to find the most likely paths of inheritance of the rare disease-causing allele, which is a simpler problem than assigning (not necessarily rare) haplotypes across the genome. In AGDB and the associated software `PedHunter`, the problem of finding the best paths of inheritance to explain the inheritance of a rare allele is solved as a combinatorial optimization problem closely related to the classical minimum Steiner tree problem in graphs [59] and large instances can be solved to optimality using mixed integer linear programming (e.g., [60]) even though the Steiner tree problem is NP-complete. In the BALSAC http://balsac.uqac.ca/ project that contains a large genealogy of the population of Québec, the rare allele inheritance problem is formulated statistically and is solved in the software ISGen https://github.com/DomNelson/ISGen using Monte Carlo Markov Chain techniques [61] extending earlier seminal work of Geyer and Thompson on a Hutterite genealogy [62]. For the haplotype problem, the methods in `thread` are based on deterministic combinatorial algorithms, like the methods in `PedHunter`. One could imagine instead solving the haplotype assignment problem by MCMC methods.

Individual-level reconstruction opens the door for many types of downstream analysis. Using reconstructed genomes to augment GWAS could increase sample sizes by hundreds of individuals when the phenotype is known. More generally, quantifying allele frequency changes, transmission distortion, and un-reconstructable (“lost”) regions of the genome allows us to model genome dynamics on a recent time scale. `thread` could be applied to other genetically characterized endogamous populations with high levels of recessive traits, such as Mennonites and Hutterites [63]. Our method would also be suitable for model organisms and domestic animals, where extensive pedigree records are common.

Our results could also be used to find individuals of clinical significance in cases where a gene-inhibiting drug may provide a therapeutic option for a disease. More specifically, loss of function (LoF) mutations in some genes have shown to protect against disease [64, 65]. If a loss of function allele has been observed, then haplotype reconstruction in any large pedigree makes it possible to identify carriers. If in addition, the pedigree has one or more loops, then haplotype reconstruction makes it possible to identify extremely rare individuals who may be homozygous for the LoF allele.

## Conclusion

In this work, we gave a formal algorithmic treatment of the problem of reconstructing ancestral haplotypes from the genotypes of individuals in the most recent generations of an extended pedigree. In publicly available software called `thread`, we designed and implemented a new algorithm for ancestral haplotype reconstruction that can handle highly complex pedigrees, which arise in isolated human populations and in animal breeding. We evaluated the performance of our algorithm on an Old Order Amish pedigree of 1338 individuals, many of whom have been diagnosed with bipolar or other mood disorders. Versions of this pedigree have been studied in psychiatric and statistical genetics for decades, as the genetic variations contributing to bipolar disorder remain poorly understood. Using our new algorithm, it is possible to trace the inheritance of many haplotypes in this pedigree, including those of long-deceased individuals. In future clinical studies, this should lead to a better understanding of which haplotypes are associated with bipolar disorder in this Amish pedigree. We anticipate that `thread` could be similarity useful in other endogamous populations.

## Supporting information

See attached document “Supplementary Material” for more information.

**S1 Fig. Pedigree structure: 1338 individuals over 10 generations**. Squares represent males and circles represent females. Dotted lines connect the same individual appearing in two different parts of the pedigree. Filled in symbols represent genotyped individuals.

**S2 Fig. Inbreeding coefficients for each individual in the pedigree structure from Figure S1**. Inbreeding coefficients were computed using the software `PedHunter` and the entire AGDB.

**S3 Fig. Simulation results per individual**. The generation of the individual is plotted on the *x*-axis (higher number generations are more recent). For the chromosomes we were able to reconstruct, we compute and plot the accuracy on the *y*-axis. Symbol size and color shows how many chromosomes we were able to reconstruct.

**S4 Fig. Demonstration of removed segments in simulations**. Headers for each set of figures show the individual ID and overall reconstruction accuracy for chromosome 21. Each horizontal line represents one IBD segment shared with a cohort of genotyped individuals. The top row of figures for each individual represent IBDs assigned at the beginning of each iteration. The bottom row represents the IBDs that remain after conflict resolution. Light blue segments are correct segments added during iteration 0, and dark blue segments are correct and added during iteration 1. Gray segments are incorrect. In the top individual, we successfully removed these gray segments and achieved 100% reconstruction accuracy. In the second individual, we failed to detect several incorrect IBD segments and achieved a lower reconstruction accuracy.

**S5 Fig. Position of reconstructed individuals in the pedigree**. Black: genotyped individual, white: no genotyped descendants, yellow-red heatmap: represents number of chromosomes reconstructed, blue: no chromosomes reconstructed.

**S6 Fig. Unsuccessful reconstruction examples**. Each horizontal line represents one IBD segment shared with a cohort of other genotyped individuals. IBD segments of the same color represent haplotypes, and have a consistent sequence along the chromosome. Small vertical lines represent heterozygous sites between the two haplotypes. A) Occasionally we only build one haplotype (which may not actually be unsuccessful if the individual was entirely homozygous for the given chromosome). B) Sometimes we have a fairly strong reconstruction, but due to the presence of other groups it does not meet our threshold for two strong group. C) Four groups may indicate ambiguity with a spouse or other close relative. D) Sometimes we see many groups and cannot resolve the individual.

**S7 Fig. Source and path distributions for chromosome 21**. (left) Distribution of the number of potential sources per IBD segment. (right) Number of paths per source (truncated at 1000, but there is an extremely long tail).

**S1 Table. Algorithm experimentation: simulated data**. The first block of results shows the output of `thread` when run on the true IBD segments from `Ped-sim`. The second block of results shows setting A of the strong groups criteria, which requires more IBD support for individuals in ancient generations (described in more detail the main text). Similarly, the third block of results shows setting B, which requires more IBD support but relaxes the length requirements for older generations. The third column shows the number of unique IBDs (called by `GERMLINE` in the second two blocks). The fourth column shows how many iterations the algorithm needed to converge. The fifth column shows the number of ancestral (ungenotyped) individuals we were able to successfully reconstruct. The last column shows the average sequence similarity of the individuals we were able to reconstruct, as compared to their true genomes.

## Acknowledgments

The authors would like to thank Jeff Knerr and Joe Cammisa for providing invaluable computational support, Amy Williams for guidance with simulations, Iain Mathieson for feedback on the method and manuscript, and the participants in the Amish Study of Major Affective Disorder (ASMAD). This research is supported in part by the National Institutes of Health, National Cancer Institute. This work utilized the computational resources of the NIH HPC Biowulf cluster http://hpc.nih.gov. YBS is funded by NIH grant no. 1F32HG011202-01. KF is funded by a Robbins/Chang Summer Fellowship. This study was supported by the NIH grant R01MH093415.

## Author Contributions

Conceptualization: RLK, MB, SM. Data Curation: AAS, RLK. Formal Analysis: KF, MK, GB, TD, ST, SM. Methodology: KF, MK, GB, YBS, AAS, RLK, MB, SM. Resources: SR, AAS. Software: KF, MK, GB, TD, ST, SM. Supervision: MB. Validation: YBS, AAS, TD, ST, SM. Visualization: KF, SM. Writing – Original Draft Preparation: KF, SM. Writing – Review & Editing: KF, GB, YBS, AAS, RLK, MB, SM.

