## Supplementary Material for "Ancestral Haplotype Reconstruction in Endogamous Populations using Identity-By-Descent"

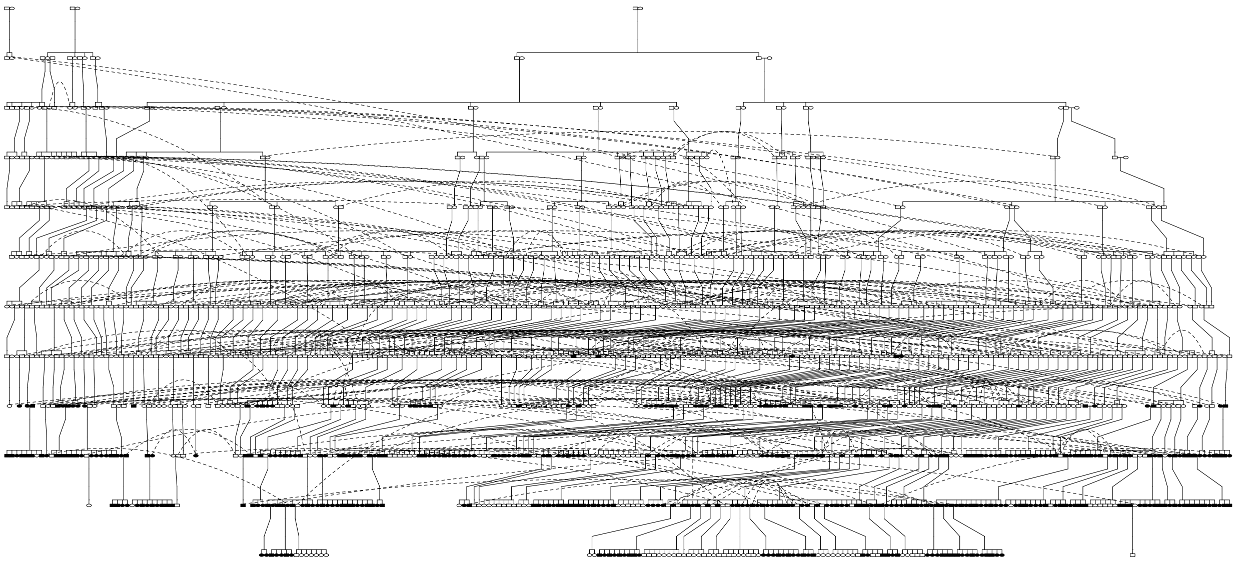

Figure S1: *Pedigree structure: 1338 individuals over 10 generations. Squares represent males and circles represent females. Dotted lines connect the same individual appearing in two different parts of the pedigree. Filled in symbols represent genotyped individuals.*

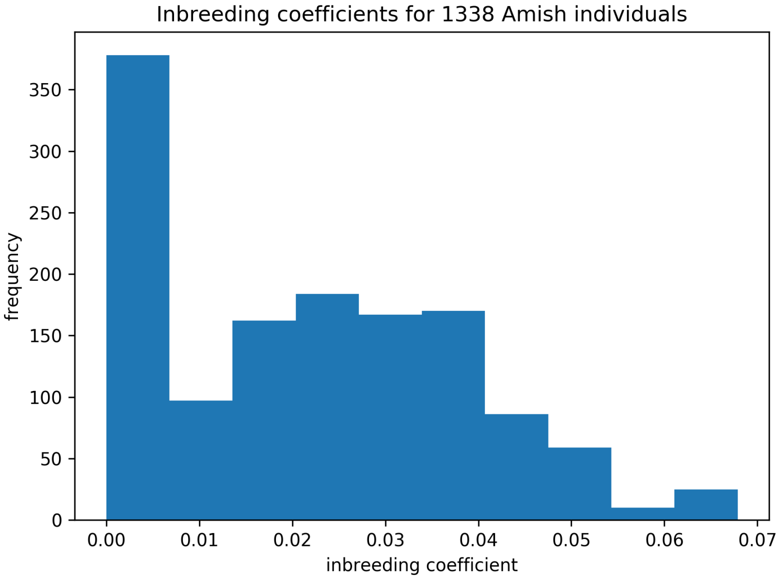


Figure S2: *Inbreeding coefficients for each individual in the pedigree structure from Figure S1. Inbreeding coefficients were computed using the software* **PedHunter** *and the entire AGDB.*

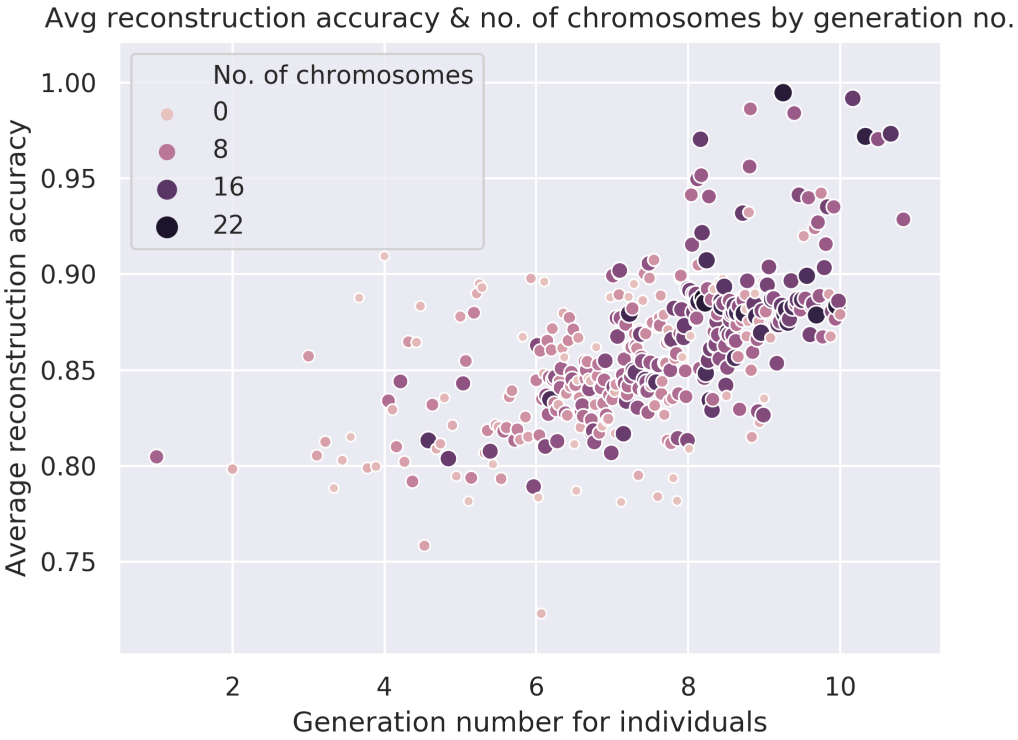


Figure S3: *Simulation results per individual. The generation of the individual is plotted on the x-axis (higher number generations are more recent). For the chromosomes we were able to reconstruct, we compute and plot the accuracy on the y-axis. Symbol size and color shows how many chromosomes we were able to reconstruct.*

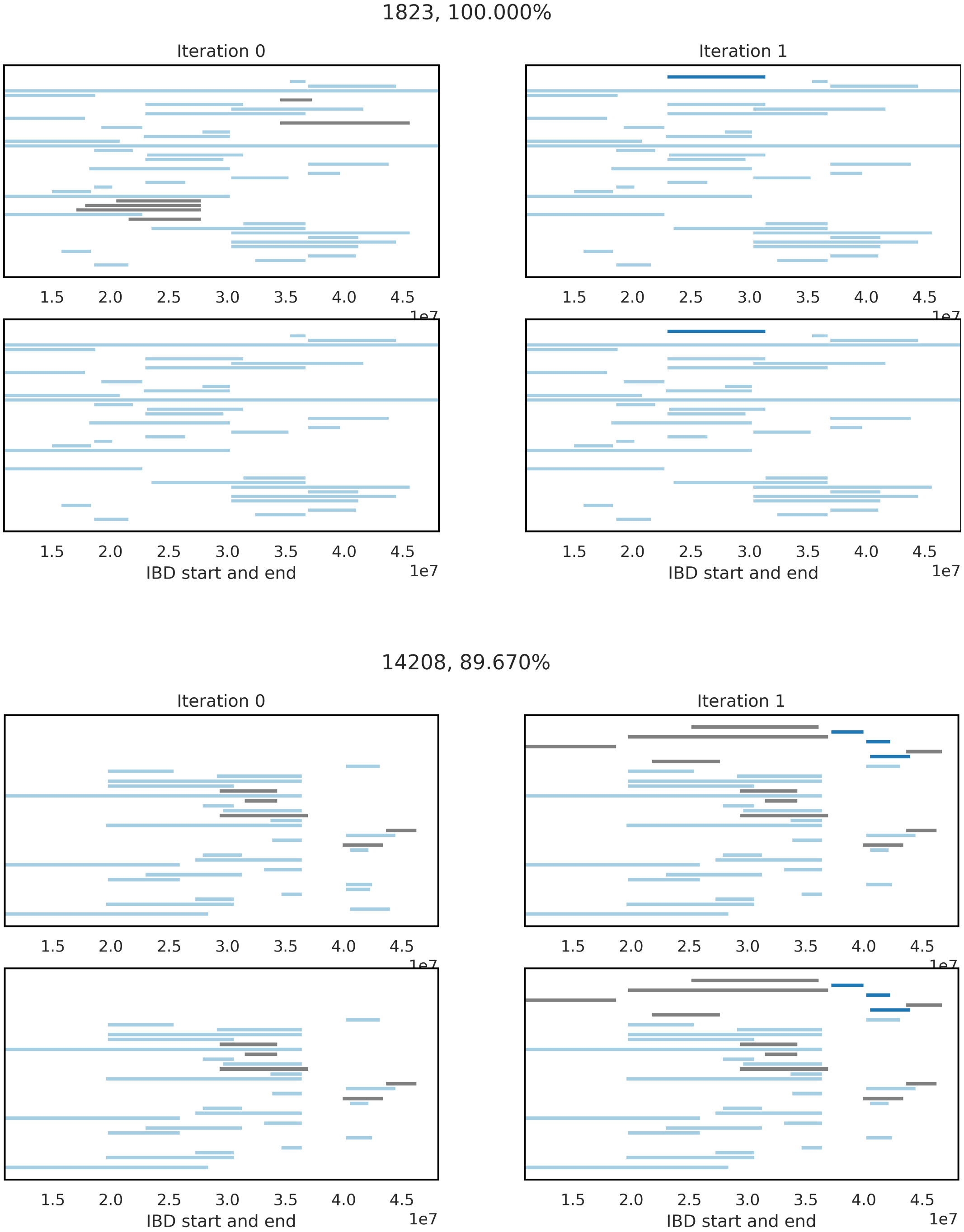


Figure S4: *Demonstration of removed segments in simulations. Headers for each set of figures show the individual ID and overall reconstruction accuracy for chromosome 21. Each horizontal line represents one IBD segment shared with a cohort of genotyped individuals. The top row of figures for each individual represent IBDs assigned at the beginning of each iteration. The bottom row represents the IBDs that remain after conflict resolution. Light blue segments are correct segments added during iteration 0, and dark blue segments are correct and added during iteration 1. Gray segments are incorrect. In the top individual, we successfully removed these gray segments and achieved 100% reconstruction accuracy. In the second individual, we failed to detect several incorrect IBD segments and achieved a lower reconstruction accuracy.*

| setting | chrom | # unique IBDs | # iter | # reconstructed | accuracy % |
|---|---|---|---|---|---|
| `Ped-sim` IBDs | 18 | 3833 | 5 | 92 | 86.66 |
| | 19 | 4109 | 5 | 154 | 86.96 |
| | 20 | 4223 | 5 | 130 | 87.79 |
| | 21 | 2367 | 6 | 97 | 88.19 |
| | 22 | 2780 | 3 | 70 | 88.78 |
| strong groups: | 18 | 3885 | 4 | 114 | 87.75 |
| setting A | 19 | 3523 | 6 | 110 | 87.17 |
| | 20 | 3959 | 5 | 106 | 87.59 |
| | 21 | 2109 | 8 | 69 | 89.87 |
| | 22 | 2528 | 4 | 65 | 87.14 |
| strong groups: | 18 | 3885 | 7 | 155 | 86.83 |
| setting B | 19 | 3523 | 4 | 125 | 86.83 |
| | 20 | 3959 | 6 | 137 | 87.17 |
| | 21 | 2109 | 4 | 88 | 88.89 |
| | 22 | 2528 | 5 | 89 | 86.73 |

Table S1: *Algorithm experimentation: simulated data. The first block of results shows the output of* ***`thread`*** *when run on the true IBD segments from* ***`Ped-sim`***. *The second block of results shows setting A of the strong groups criteria, which requires more IBD support for individuals in ancient generations (described in more detail the main text). Similarly, the third block of results shows setting B, which requires more IBD support but relaxes the length requirements for older generations. The third column shows the number of unique IBDs (called by* ***`GERMLINE`*** *in the second two blocks). The fourth column shows how many iterations the algorithm needed to converge. The fifth column shows the number of ancestral (ungenotyped) individuals we were able to successfully reconstruct. The last column shows the average sequence similarity of the individuals we were able to reconstruct, as compared to their true genomes.*

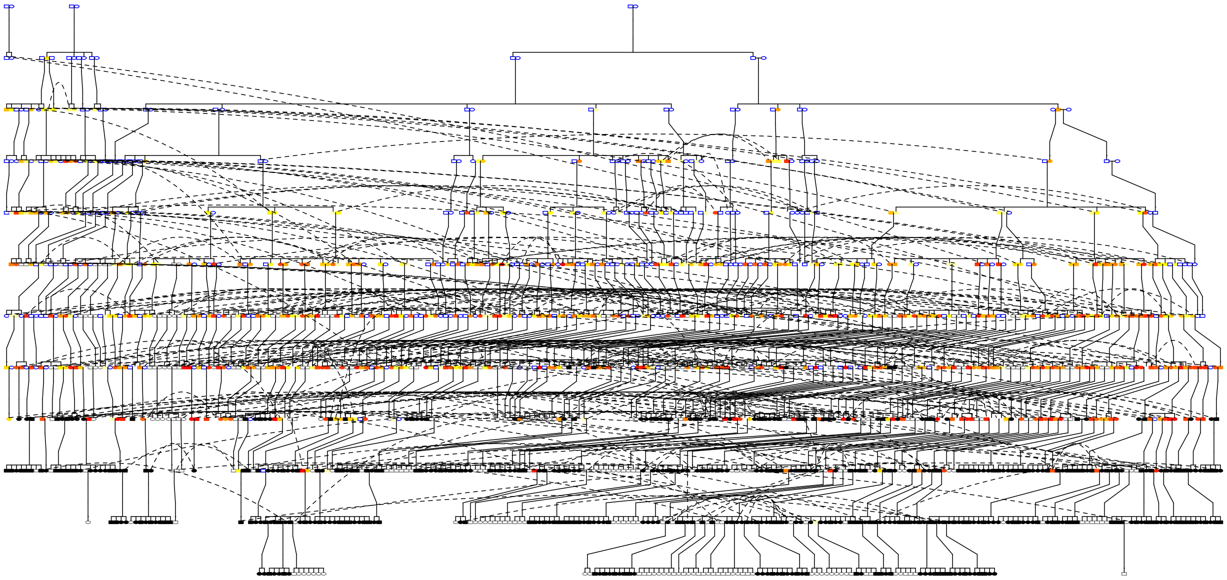

Figure S5: *Position of reconstructed individuals in the pedigree: colors are as follows. Black: genotyped individual, white: no genotyped descendants, yellow-red heatmap: represents number of chromosomes reconstructed, blue: no chromosomes reconstructed.*

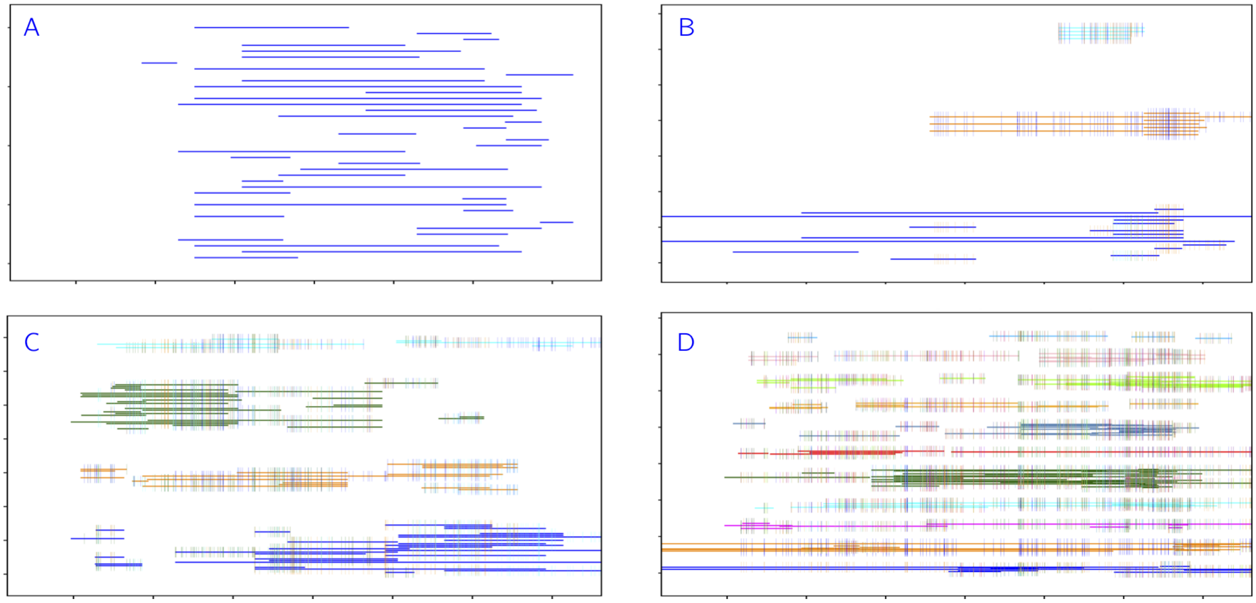


Figure S6: *Unsuccessful reconstruction examples: Each horizontal line represents one IBD segment shared with a cohort of genotyped individuals. IBD segments of the same color represent haplotypes, and have a consistent sequence along the chromosome. Small vertical lines represent heterozygous sites between the two haplotypes. A) Occasionally we only build one haplotype (which may not actually be unsuccessful if the individual was entirely homozygous for the given chromosome). B) Sometimes we have a fairly strong reconstruction, but due to the presence of other groups it does not meet our threshold for two strong group. C) Four groups may indicate ambiguity with a spouse or other close relative. D) Sometimes we see many groups and cannot resolve the individual.*

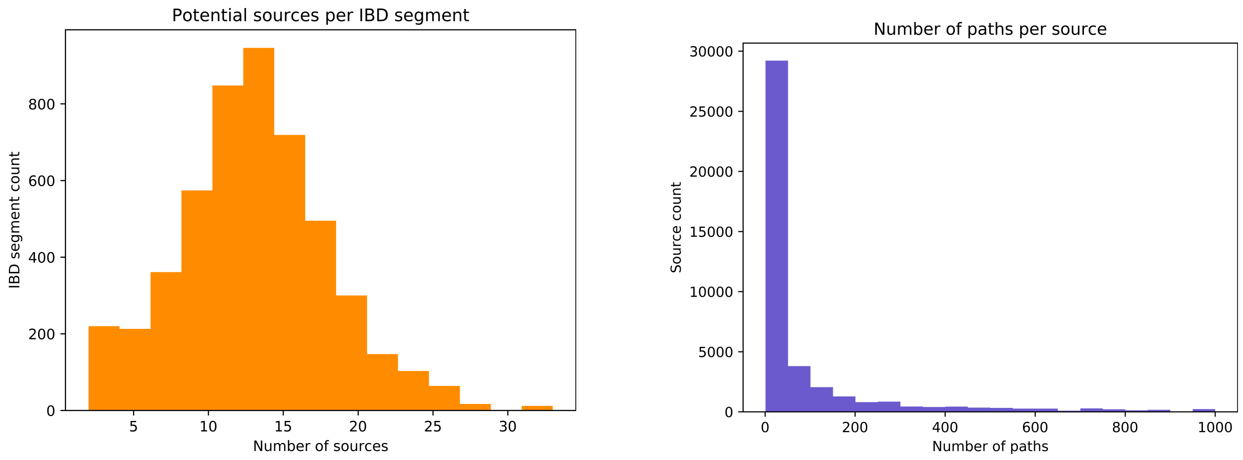


Figure S7: *Source and path distributions for chromosome 21. (left) Distribution of the number of potential sources per IBD segment. (right) Number of paths per source (truncated at 1000, but there is an extremely long tail).*

**Algorithm 1:** Overview

**Input:** $G$ = genotyped individuals, $NG$ = non-genotyped individuals, $\mathcal{P}$ = pedigree tree relating all individuals in $G$ and $NG$

**Output:** $R$ = reconstructed individuals, $\mathcal{G}_p$ = groups for each individual $p \in R$

find IBDs shared between $G$ using `GERMLINE`

**for** $I_k \in IBDs$ **do**
- $C_k$ = cohort of individuals from $G$ sharing $I_k$
- $S_k$ = sources of $C_k$ (Algorithm 2)
- $d_k(s)$ = number of descendance paths for each $s \in S_k$ (Algorithm 2)

**end**

$R = G$

$IS$ = list of IBDs to source

**while** *$R$ not changing and $IS$ not empty* `// this loop tracks iterations` **do**
- **for** $I_k \in IS$ **do**
  - **while** *assignment unsuccessful and $S_k$ is not empty* **do**
    - selected source $s^* = \arg\min_s d_k(s)$
    - **if** $d_k(s^*) >$ *path threshold* **then**
      - ignore $I_k$
    - **end**
    - **else**
      - individuals $D_k(s^*)$ = all individuals lying on each path from $s^*$ to $C_k$
      - assign $I_k$ to all individuals in $D_k(s^*)$
      - **if** *$I_k$ conflicts with reconstructed individual in $D_k(s^*)$* **then**
        - remove $I_k$ from all $D_k(s^*)$
        - remove $s^*$ from $S_k$
        - assignment is unsuccessful
      - **end**
    - **end**
  - **end**
- **end**
- reset $IS$ to empty list
- **for** *individual $p \in NG$* **do**
  - $\mathcal{G}_p$ = reconstructed haplotype groups (Algorithm 3)
  - **if** *exactly 2 strong groups in $\mathcal{G}_p$* **then**
    - add $p$ to $R$
  - **end**
  - **if** *2 strong groups and one or more weak groups in $\mathcal{G}_p$* **then**
    - remove weak groups from $\mathcal{G}_p$
    - add all IBDs from weak groups to $IS$
    - add $p$ to $R$
  - **end**
- **end**

**end**

**return** *$R$, $\mathcal{G}_p$ for each $p \in R$*

**Algorithm 2:** Source and Descendance Path Finding

**Input:** $C$ = a cohort of individuals sharing a single IBD, $\mathcal{P}$ = pedigree tree containing relationships between individuals

**Output:** $S$ = a list of possible non-redundant sources for cohort $C$

```
queue Q = list(C)
for cohort member p ∈ C do
    multiset M_p = {p}
end
while Q is not empty do
    individual p = Q.pop
    if p is married-in then
        skip the following (married-in have no known ancestors)
    end
    if p^(f) has not been processed then
        father's multiset M_f = M_p
        father's children set CH_f = p
        add father to Q
    end
    else
        extend father's multiset M_f by M_p
        add p to father's children set CH_f
        add M_p and p to M and CH of any processed ancestors of father
    end
    repeat process for p^(m)
end
sources S = all individuals p s.t. M_p contains all c ∈ C
for source s ∈ S do
    M_chmax = largest M_ch for ch ∈ CH_s
    if length of M_s = M_chmax then
        remove redundant source s from S
    end
end
for source s ∈ S do
    if s.spouse in S and M_s = M_s.spouse then
        remove s and s.spouse from S
        add couple s&s.spouse to S, s.t. M = M_s and CH = CH_s
    end
end
for source s ∈ S do
    number of descendance paths d(s) = ∏_{c∈C} m_s(c), where m_s(c) = multiplicity of c in M_s
end
return S and d(s) for all s ∈ S
```

**Algorithm 3:** Grouping

**Input:** $R$ = genotyped or reconstructed individuals, $A$ = non-reconstructed individuals, ungrouped IBDs $\mathcal{I}_p$ have been placed in each individual $p$

**Output:** $\mathcal{G}_p$ = groups for each individual $p$

```
for individual p ∈ R do
    for IBD I ∈ I_p do
        add I to one or both groups in G_p depending on zygosity
    end
end
for individual p ∈ A do
    find any homozygous groups G_p^(o)
    use overlapping IBDs in I_p to build heterozygous groups G_p^(e)
    duplicate groups in G_p^(o) and create G_p = G_p^(o) ∪ G_p^(e)
    remove all IBDs from S_p that were used to build groups in G_p
    for pairs of groups G_i, G_j ∈ G_p and remaining IBD I ∈ I_p do
        if I overlaps G_i and G_j sufficently then
            merge G_j into G_i and delete G_j
        end
    end
    for pairs of groups G_i, G_j ∈ G_p do
        if G_i and G_j overlap or "line up" then
            merge G_j into G_i and delete G_j
        end
    end
end
```

### Complexity Analysis

The worst-case time complexity analysis involves several incomparable quantities. We use:

$$
\begin{aligned}
|G| &= \text{number of genotyped individuals} \\
|NG| &= \text{number of ungenotyped individuals with a genotyped descendant} \\
|R| &= \text{number of reconstructed individuals} \\
|S| &= \text{number of sources} \\
|P| &= \text{number of individuals in the pedigree} \\
|A| &= \text{number of generations}s
\end{aligned}
$$

and note that $|G| \leq |P|$, $|NG| \leq |P|$, $|R| \leq |P|$, and $|S| \leq |P|$. Additionally, we use:

$$
\begin{aligned}
|I| &= \text{number of unique IBD segments} \\
I_{\max} &= \text{maximum number of SNPs within an IBD segment} \\
|H| &= \text{total number of SNPs on the chromosome} \\
|J| &= \text{sum of number of SNPs in all IBDs, which is bounded by} I_{\max} \cdot |I| \\
|C| &= \text{sum of all cohort sizes} \\
r &= \text{number of iterations of the algorithm}
\end{aligned}
$$

We analyze four parts:

1. processing of `GERMLINE` output as input to `thread`
2. Algorithm 3
3. Algorithm 2
4. Algorithm 1

Each row of `GERMLINE` output reports two individuals sharing one IBD. Therefore the size of the `GERMLINE` output file is $O(|I||G|^2)$, which we simplify to $O(|I||P|^2)$. These rows have to be processed to assign to each individual their corresponding IBDs and compile lists of individuals sharing each IBD. The two gathering tasks can be implemented to take time $O(|I||P|^2)$ and in a memory efficient manner if we assume that the `GERMLINE` output is sorted so that all rows for the same IBD are consecutive. However, the current implementation takes time $O(|I||P^3|)$ because it uses a general and inefficient implementation of union-find to determine the set of individuals carrying each IBD. In this context, the sets are finite and we know that the final result will be a single set for each IBD, so union-find can be implemented in linear $O(|P|)$ time using bit arrays.

The time-consuming steps in Algorithm 3 compare IBDs for overlap and conflicts. In the worst case one might have to compare every SNP in one IBD to every SNP in another IBD. Algorithm 3 is run for every ungenotyped individual, so the worst case running time for all uses of Algorithm 3 in one iteration is $O(|NG||J|^2)$, which we simplify to $O(|J|^2|P|)$.

In Algorithm 2, building the multisets takes $O(|P|^2)$ time. Testing for sources takes $O(|P||C|)$ time. Removing redundant sources takes $O(|P|)$ time. Replacing single sources by couples takes $O(|P|)$ time. Computing products of the number of paths takes $O(|S||C|)$ time, which is $O(|P||C|)$. The dominant time is to build all the paths from one source to one target, which $O(|P|^2 \cdot 2^{|A|})$, where $A$ is the number of generations. Overall, this is

$$O\left(|P|^2 \cdot 2^{|A|}) + |P||C|\right)$$

since $|S| < |P|$.

There are three time-consuming parts of Algorithm 1 with different complexities. The assignment steps require $O(I_{\max}|I||P|)$ time. The costs of identifying conflicts and overlaps to make the groups for each individual is $O(|H||P|)$ with a careful implementation of these two tests. As indicated above the cost of all the calls to Algorithm 3 is $O(|J|^2|P|)$. Putting the three terms together, we get $O(I_{\max}|I| + |H| + |J|^2)|P|$. For this data set and any interesting endogamous data set, we would expect the $O(|J|^2|P|)$ term to dominate.

Since the cost of Algorithm 3 is subsumed in Algorithm 1, we do not include it in the final tally. Algorithms 1 and 2 are used for $r$ iterations, while the processing of `GERMLINE` output is done only one time. Putting together the costs of preprocessing and Algorithms 1 and 2, we arrive at a complexity of

$$O\left(|I||P|^3 + r(|P|^2 \cdot 2^{|A|} + |C||P| + I_{\max}|I| + |H| + |J|^2|P|)\right)$$

In a typical dataset, we would expect $|P|$ to be far smaller than $|H|$, $|I|$, or $|J|$ and that is true for this dataset. Thus, we expect that the dominant would be $O(r|J|^2|P|)$ or the incomparable exponential term $O(r|P|^2 \cdot 2^{|A|})$. Since these two terms are incomparable, we performed runtime profiling with the `python` module `cProfile`. These experiments validated that the procedures taking $O(r|J|^2|P|)$ and $O(r(|P|^2) \cdot (2^{|A|}))$ dominate. For the pedigree structures tested, on the shortest chromosomes (21, 22), these two parts of `thread` take similar amounts of time, within a multiplicative factor of 2. On most of the (longer) chromosomes the $O(r|J|^2|P|)$ term dominates because $|J|$ increases with the length of the chromosome, but the costs of finding paths from sources $O(r|P|^2 \cdot 2^{|A|})$ does not depend substantially on the chromosome length; on the longer chromosomes, one might need to consider more potential sources, but this consideration does not affect the asymptotic upper bound on running time.

### Probabilistic source identification

Given an IBD segment $I$ and associated cohort $C$ (genotyped individuals that carry $I$), we wish to approximate the probability that each potential source $(s_1, s_2, \cdots, s_k)$ is the true origin of $I$. At a high level, we compute this by compiling the probabilities of transmitting the IBD from the source to each member of the cohort.

Let the genetic distance of $I$ be $d$ cM. Then the probability of a recombination event within the IBD segment during one meiosis is:

$$r = \frac{1 - e^{-2d/100}}{2}$$

First we will compute the probability that a child receives 0, 1, or 2 copies of the IBD from its parents. Let $[f_0, f_1, f_2]$ be the probabilities that the first parent has 0, 1, or 2 copies of the IBD, and similarly for the second parent $[m_0, m_1, m_2]$. Then we can compute the child probabilities as follows:

$$
\begin{aligned}
c_0 &= \left(f_0 + \frac{1}{2} f_1 (1+r)\right)\left(m_0 + \frac{1}{2} m_1 (1+r)\right) \\
c_1 &= \left(f_0 + \frac{1}{2} f_1 (1+r)\right)\left(\frac{1}{2} m_1 (1-r) + m_2\right) + \left(\frac{1}{2} f_1 (1-r) + f_2\right)\left(m_0 + \frac{1}{2} m_1 (1+r)\right) \\
c_2 &= \left(\frac{1}{2} f_1 (1-r) + f_2\right)\left(\frac{1}{2} m_1 (1-r) + m_2\right)
\end{aligned}
$$

Then, for each source $s_i$ we assume there is one copy of the IBD to start (i.e. the probabilities are [0,1,0]). In the case of a couple source, we arbitrarily choose one parent to have the single copy of the IBD segment. We maintain a queue of individuals, which starts out with the children of the source. For each individual in the queue, we compute the probabilities of IBD transmission given their parent probabilities, and then add their children to the queue. Whenever we reach a cohort individual $a$, we retain $a_1$ if the individual had one copy and $a_2$ if they had two copies (we know the individual had at least one copy since they are in the cohort for this IBD). Finally, we compute the average of these retained probabilities, which we denote $P(s_i)$. To choose a source, we take the max:

$$s^* = \arg\max_i P(s_i)$$

Then the algorithm proceeds as follows – if this was a bad source (conflicts with genotyped or reconstructed individuals), we still remove it and choose the source with the next highest probability. Also note that these probabilities do not sum to 1, but could be normalized to do so. Finally, note that these probabilities are approximations of the complete probability for a particular configuration, as they do not include the probability the IBD segment was *not* transmitted to genotyped individuals outside the cohort.
